# Evolution of predictive memory in the hippocampus

**DOI:** 10.1101/2022.09.08.507204

**Authors:** Adam M. P. Miller, Alex D. Jacob, Adam I. Ramsaran, Mitchell L. De Snoo, Sheena A. Josselyn, Paul W. Frankland

## Abstract

The brain organizes experiences into memories that can be used to guide future behavior. Hippocampal CA1 population activity may reflect the retrieval of predictive models that contain information about future events, but little is known about how these kinds of memories develop with experience. We trained mice on a series of tone discrimination problems with or without a common statistical structure to observe how memories are formed and updated during learning. Mice that learned structured problems integrated their experiences into a predictive model that contained the solutions to upcoming novel problems. Retrieving the model during learning improved discrimination accuracy and facilitated learning by decreasing the amount of new information that needed to be acquired. Using calcium imaging to track the activity of thousands of CA1 neurons during learning on this task, we observed the emergence of a persistent hippocampal ensemble at the same time that mice formed a predictive model of their environment. This ensemble was reactivated during training and incorporated new neuronal activity patterns from each training problem. Interestingly, the degree to which mice reactivated the ensemble was related to how well their model predicted the content of the current problem, ensuring that the model was only updated with congruent information. In contrast, mice trained on unstructured problems did not form a predictive model or engage a persistent ensemble. These results show how hippocampal activity supports building predictive models by organizing newly learned information according to its congruence with existing memories.

## Introduction

All animals use information from their past to guide their present behavior. Memories derived from multiple experiences may be particularly useful for supporting efficient behavior by providing insight into the rules that govern environments. This type of memory, often described as a predictive model, is considered a fundamental feature of high-level cognitive functions such as creativity and intelligence (Kumaran et al., 2016; Tervo et al., 2016; Behrens et al., 2018; Momennejad, 2020; Morton and Preston, 2021). However, little is understood about how predictive models are formed in the brain and used to guide behavior.

The hippocampus has been implicated in supporting predictive models across several species (Zeithamova et al., 2012; Pudhiyidath et al., 2022; Brunec and Momennejad, 2022; Vikbladh et al., 2019; Schapiro et al., 2016; Knudsen and Wallis, 2021; Baraduc et al., 2019; Bulkin et al., 2016; McKenzie et al., 2014; Nieh et al., 2021; Tse et al., 2007). One possibility is that the hippocampus supports the development of predictive models by repeatedly retrieving and updating memories with new information (Lee, 2009; Schlichting and Frankland, 2017; Gisquet-Verrier and Riccio, 2018; Mack et al., 2018; Mau et al., 2020). In this framework, animals may infer a predictive model of their environment from a subset of possible observations and then interpret subsequent experiences in terms of this model (i.e. latent state theory; Fuhs and Touretzky, 2007; Gershman and Niv, 2010; Niv, 2019; Redish et al., 2007). When an existing predictive model can explain the animal’s current experience, it is retrieved along with a corresponding hippocampal activity pattern (i.e. map or context code; Sanders et al., 2020; Whittington et al., 2020). New learning that is consistent with the model may then be integrated into the model, whereas new learning that is not consistent with any existing model may prompt the formation of a new model.

Although it is challenging to study mental representations in the brain (Tervo et al., 2016), several aspects of this idea have been examined. For instance, it has been shown that the hippocampus encodes experience in state spaces (Wood et al., 2000; Nieh et al., 2021; Sun et al., 2020; Samborska et al., 2021) and that pre-existing memories (McKenzie et al., 2014; Tse et al., 2007; Cai et al., 2016) and neural structure (Epsztein et al., 2011; Dragoi and Tonegawa, 2011; Sadtler et al., 2014; McKenzie et al., 2021) can affect learning. However, whether the interplay between new learning and existing memory accrued over many experiences gives rise to a predictive model that guides behavior has not been assessed directly, especially at the neural level. Here, we developed a new behavioral protocol that repeatedly probes the task representation of a mouse as it learns a series of unique problems. At the behavioral level, we found that mice formed a predictive model that accurately predicted the solution to novel problems by integrating their memories of past problems, and that mice retrieved and updated their model with new learning only when what they learned matched their predictions. At the neural level, we used calcium imaging to track changes in hippocampal CA1 ensemble activity as mice formed and retrieved a predictive model. We found that mice formed a persistent hippocampal ensemble by incorporating neurons associated with prior training problems, and that they reactivated this ensemble during new learning when they accurately predicted the solutions to the problems.

## Results

### Mice show superior learning of structured experiences

We trained 80 mice to perform a novel auditory discrimination and foraging task to examine how memory-based predictions are formed and updated. In this task, mice hear tones and learn to wait longer in response to certain frequencies in order to obtain chocolate milk rewards (**Figure 1A**; **Supplemental figure 1**). Training was divided into a series of six unique training problems, and mice only advanced to the next problem after successfully learning the current one. Half of the mice (n=40) received a Structured training protocol where the highly rewarded (i.e., rich) tones from every problem fell within a continuous band of frequencies (**Figure 1B**), while the other mice (n = 40) received an Unstructured training protocol where the rich and poor tones were more evenly distributed (**Figure 1C**). Consistent with previous work on memory transfer (Harlow, 1949), the Structured training group showed increased preference for the rich tone on the first day of training of later problems (one-way repeated measures ANOVA, F(5, 195) = 23.99, p < 0.001; **Figure 1D**), as well as more rapid learning over the course of training (F(5, 195) = 18.45, p < 0.001; **Figure 1F**), even though every problem was novel, indicating that early training experiences enhanced the ability of mice in the Structured training group to learn novel problems. By contrast, the Unstructured training group reliably preferred the poor tone (F(5, 195) = 3.76, p < 0.01; **Figure 1E**) and often showed slowed learning (F(5, 195) = 4.59, p < 0.001; **Figure 1G**), indicating memory interference. Direct comparisons between the groups on the first and last problems (which were the same in both training protocols; i.e., the common problems) confirmed that only the Structured training group improved their performance on the first day of training problems (two-way repeated measures ANOVA, group X problem interaction, F(1, 78) = 93.89, p < 0.001; between groups on problem 1, t(78) = 0.13, p = 0.90; between groups on problem 6, t(78) = 11.43, p < 0.001; **Figure 1H**) and their rate of learning (group X problem interaction, F(1, 78) = 63.18, p < 0.001; between groups on problem 1, t(78) = 0.81, p = 0.42; between groups on problem 6, t(78) = 9.80, p < 0.001; **Figure 1I**). Lastly, the Structured training group did not require as much training to reach criterion (defined as the difference between the preference for the rich tone on the first and last day; two-way repeated measures ANOVA, group X problem interaction, F(1, 78) = 10.37, p < 0.01; between groups on problem 1, t(78) = 1.08, p = 0.28; between groups on problem 6, t(78) = 4.76, p < 0.001; Structured training group compared to zero, t(39) = 1.13, p = 0.26; Unstructured training group compared to zero, t(39) = 9.14, p < 0.001; **Figure 1J**), indicating that the last problem contained less new information for the Structured training group despite neither group having been trained on that problem before.

**Figure 1.**
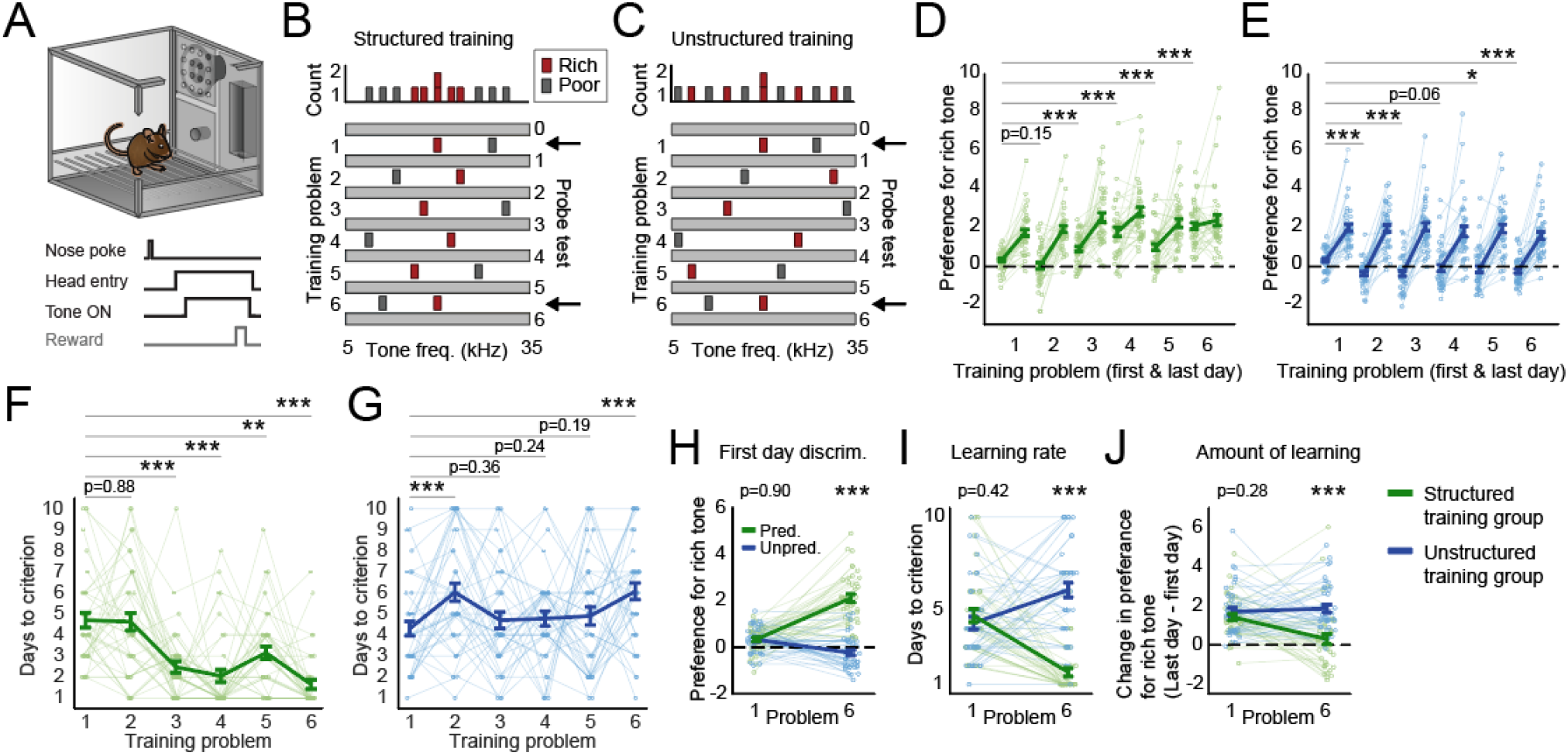
Efficient learning of structured versus unstructured problems. **(A)** Mice were trained and tested in an operant chamber with a nose-poke port, reward-delivery hopper, and a speaker that delivered different tones. Each trial consisted of a nose-poke followed by head entry into the hopper. Upon head entry, a tone played followed by a potential reward delivery. **(B)** Two groups of mice were trained on a series of tone discrimination problems. In the Structured training group, all of the rich tones fell within a continuous band of frequencies. **(C)** In the unstructured group, the rich tones were more evenly distributed. The first and last problem were the same for both groups (black arrows). **(D)** Preference for the rich tone is shown in terms of the standardized mean difference between how long mice waited in response to the rich tone and the poor tone. Preferences are shown for the first and last day of training on every problem. Mice were trained to the same criterion (longer wait times on rich tone trials than poor tone trials as determined by a significant T-test, alpha level = 0.01) on every problem and then were given one additional day of training (i.e., the last day). The Structured training group showed better discrimination on the first day of problems later in learning compared to the first problem. **(E)** The Unstructured training group did not improve first-day performance and typically showed significantly worse performance. **(F)** The number of days required to reach criterion. The Structured training group learned new problems faster later in learning. **(G)** The learning rate of the Unstructured training group failed to improve. Direct comparisons between the two groups on the first and last problem revealed that the Structured training group **(H)** showed higher initial discrimination on the last training problem, **(I)** learned the last problem in fewer days, and **(J)** learned less new information on the last problem compared to the Unstructured training group.

### Mice integrate structured experiences to form a stable, generalized model of the environment

We hypothesized that the Structured training group performed better on later discrimination problems by integrating its early experiences into a predictive model of the environment that supported new learning (Tolman, 1948; Tse et al., 2007). If this is true, then the Structured training group should develop tone preferences (increased wait times) that match the statistics of the full Structured training set (and not simply the most recent problem; **Figure 1B**). Furthermore, if this process depends on updating a predictive model with new information, then tone preferences should change (be updated) when mice encounter unpredicted problems but should remain stable when the mice encounter problems that are already predicted by the model (and therefore do not require additional learning).

To observe how tone preferences changed during Structured or Unstructured training, we examined the behavioral responses of both groups during probe tests administered before and after each training problem (**Figure 1B, C**). During probe tests, 41 unique tones (min frequency = 5kHz, max frequency = 35kHz; all tones, p(reward) = 0.5) were presented in random order to examine the length of time mice would wait in response to a range of tones, including tones that had not been trained yet. Early in training, both groups generalized beyond the two most recent training tones, preferring all tones beyond the recent rich tone to all tones beyond the recent poor tone (e.g., probes 1 and 2; see Purtle, 1973). Later in training, the Structured training group began to selectively prefer tones in the center of the auditory range (including novel tones that were not previously trained), consistent with the statistics of the training set (**Figure 2A, B**). This suggests that the Structured training group integrated its training experiences into an accurate model of the relationship between tone and reward.

**Figure 2.**
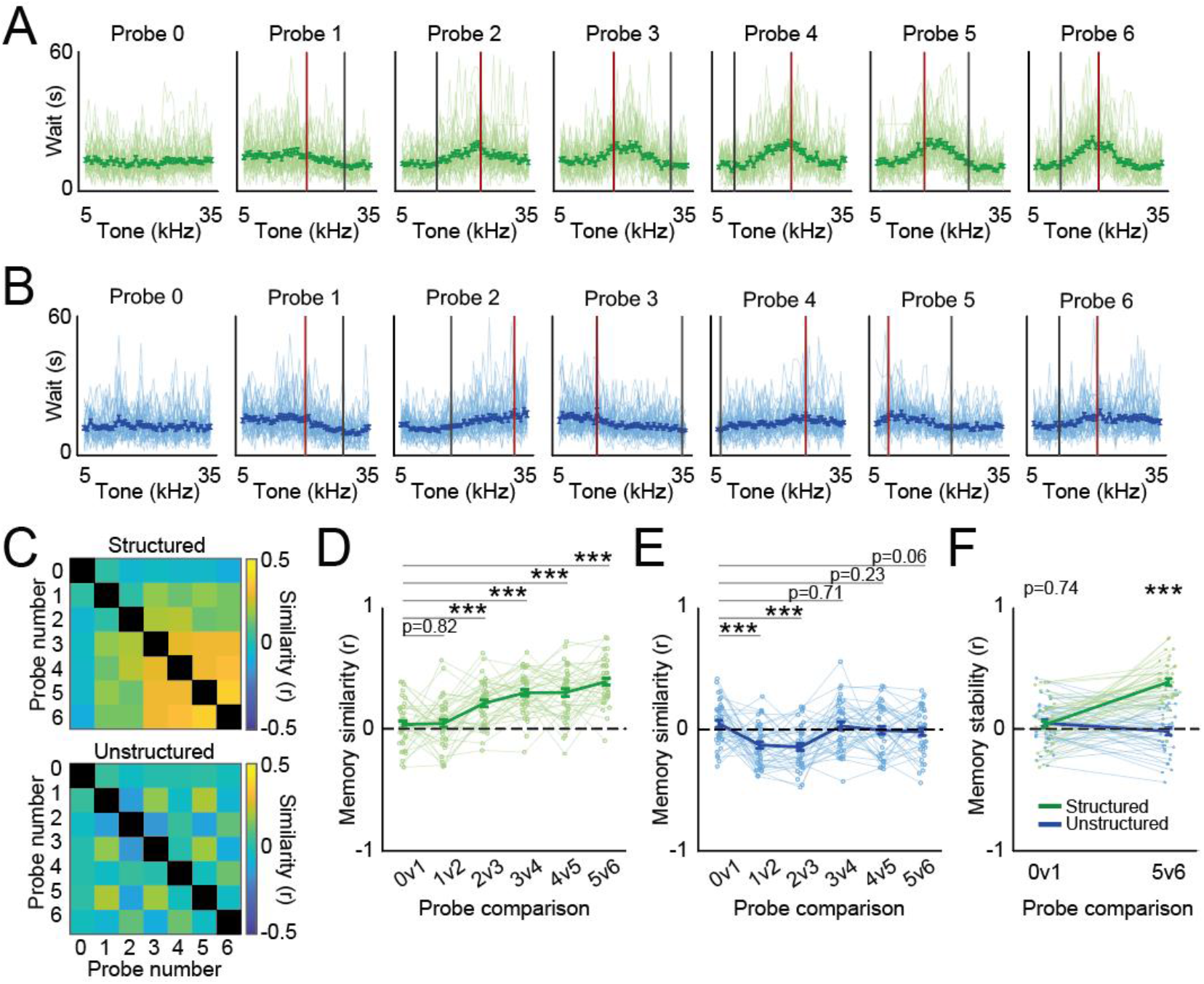
Integrating structured experiences into a generalized model. **(A)** The wait time responses to every tone are shown for every subject in every probe test of the Structured training group. Probe 0 is the pre-probe administered before any tone training. Subsequent probes were administered the day after completing the corresponding problem. The rich and poor tone and poor tone frequencies of the prior problem are shown as red and grey lines, respectively. **(B)** Same as A, but for the Unstructured training group showing no integration of new information into previous memories. **(C)** Correlation matrices showing the mean of all subject correlations between every pair of probe tests for the Unstructured training group (top) and Unstructured training group (bottom). **(D)** The correlation between every pair of sequential probe tests is shown for the Structured training group. **(E)** Same as D, but for the Unstructured training group. **(F)** The correlation between the probes before and after the two common problems (problem 1 and problem 6) are shown for both groups.

To quantify the degree to which the model changed after each training experience, we assessed the similarity of tone preferences before and after each problem. The Structured training group developed stable tone preferences during the second half of training (linear mixed-effects (LME) model with fixed effects for first probe, second probe, and group, and a random effect for subject, first probe X second probe X group interaction, t(3352) = 9.16, p < 0.001; **Figure 2C**; the Structured training group, one-way repeated measures ANOVA, F(5, 195) = 27.79, p < 0.001; **Figure 2D)**. In contrast, the tone preferences of the Unstructured training group were either uncorrelated or anticorrelated (F(5, 195) = 7.27, p < 0.001; **Figure 2E**), indicating that this group did not form a stable model. Between-group comparisons confirmed that the Structured training group developed a stable model while the Unstructured training group continued to show learning-related changes (two-way repeated measures ANOVA, group X stage interaction, F(1, 78) = 61.86, p < 0.001; Structured training group compared to zero, t(39) = 13.78, p < 0.001; Unstructured training group compared to zero, t(39) = 0.67, p = 0.50; **Figure 2F**).

### Generalized model predicts future discriminations and improves learning

The above observations indicate that the Structured training group formed a stable model at the same time that they began to show efficient learning of novel discriminations. This raises the possibility that the model (integrated memories of past problems) contained information about future (not-yet trained) problems that facilitated learning on these problems. To quantify the information about discrimination problems contained in the model, we fit a curve to the tone preferences (wait times) observed during every probe test (see methods) and then measured the difference between how long the mouse waited in response to the rich and poor training tones used in the training problems. As a proof of concept, we first asked whether the model expressed during probe tests correctly discriminated between the rich and poor tones of the most recent training problem. Because both groups successfully learned to prefer the rich tone to the poor tone over the course of every training problem (**Figure 1 D-E**), we anticipated that the mice would similarly discriminate between these tones during the subsequent probe test one day later. Consistent with this, mice showed a similar preference for the rich tone during training problems and the probe tests the following day (**Figure 3A;** every Structured training group session compared to zero, p < 0.001; **Figure 3B**; every Unstructured training group session compared to zero, p < 0.001; **Figure 3C**; direct comparisons after the common problems, two-way repeated measures ANOVA, group X probe interaction, F(1, 78) = 4.35, p = 0.04; at probe 1, t(78) = 1.14, p = 0.26; at probe 6, t(78) = 1.66, p = 0.10; **Figure 3D**), indicating that the model expressed during the probe test contained information about the discrimination learned during the preceding training problem.

**Figure 3.**
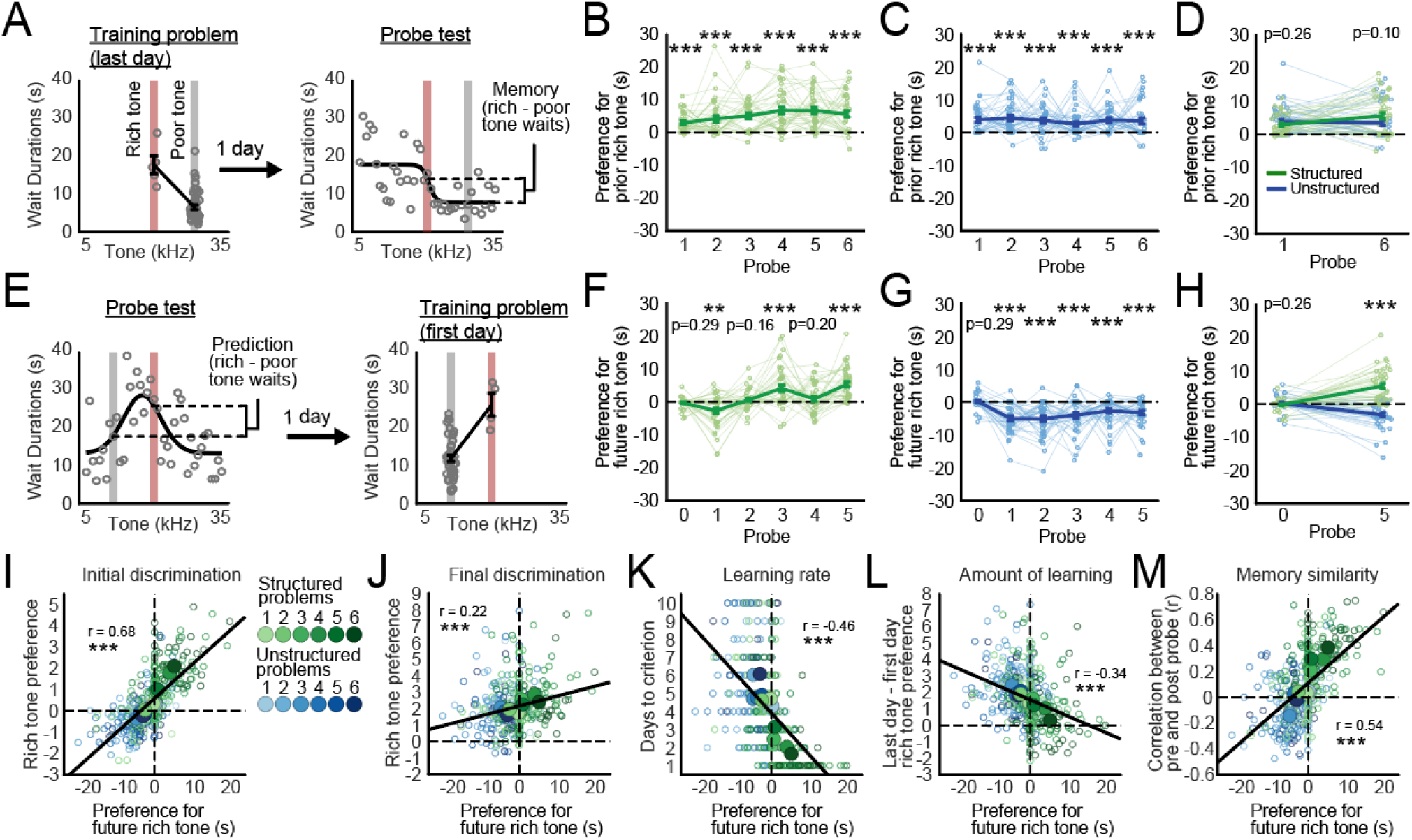
Predictive models guide new learning. **(A)** Memory for the reward values of the rich and poor tones used in the prior problem can be measured during the probe test one day after training. All wait times from two example sessions (the last day of a training problem and the probe test one day later) are shown from one mouse. Grey circles indicate the wait times on individual trials. We quantified the memory for the prior problem in terms of the difference between the how long the mouse waited in response to the same tones during the subsequent probe test. Wait times were determined by fitting a curve to the observed waits from every trial. **(B)** The Structured training group and **(C)** the Unstructured training group showed significant memory for the prior problem during every probe test. **(D)** The two groups also showed similar memory for each of the two common training problems. **(E)** Predictions for the reward values of future rich and poor tones can be measured during the probe test one day before the start of training. We measured predictions in the same way that we measured memory, but with the rich and poor tones from the upcoming (and not the prior) problem. **(F)** The Structured training group accurately predicted the solutions to some problems later in learning after initially making inaccurate predictions. **(G)** The Unstructured training group showed inaccurate predictions throughout training. **(H)** The Structured training group developed superior predictions with training compared to the Unstructured training group. **(I)** The more accurate the prediction (measured during the preceding probe test), the stronger the preference for the rich tone on the first day of training, defined as the standardized mean difference between the wait times in response to the rich and poor tones. Large circles show condition means. **(J)** The more accurate the prediction, the stronger the preference for the rich tone on the last day of training. **(K)** The more accurate the prediction, the faster the subject learned the training problem. **(L)** The more accurate the prediction, the less new information was learned during a training problem, defined as a smaller change in the preference for the rich tone from the first to the last day of training. **(M)** More accurate predictions were associated with smaller learning-related changes to the memory, defined in terms of the correlation between the behavioral responses on probe tests before and after training (higher correlations indicate smaller changes).

To examine whether the model expressed during probe tests also contained information about *future* problems, we asked whether mice waited longer in response to the upcoming rich tone than to the upcoming poor tone. Similar to above, we found that the amount of time mice waited in response to the training tones on the first day of a problem closely matched how long they waited in response to those same tones on the probe test one day prior (**Figure 3E**), indicating that the mice were retrieving the same model during training that they retrieved during the prior probe. However, only the Structured training group developed a preference for future rich tones over future poor tones (all compared to zero: probe 0, t(39) = 1.07, p = 0.29; probe 1, t(39) = 3.32, p < 0.01; probe 2, t(39) = 1.42, p = 0.16; probe 3, t(39) = 4.21, p < 0.001; probe 4, t(39) = 1.31, p = 0.20; probe 5, t(39) = 6.87, p < 0.001, **Figure 3F**). The Unstructured training group never predicted upcoming problems accurately and was often inaccurate (all compared to zero: probe 0, t(39) = 0.62, p = 0.54; probe 1, t(39) = 7.24, p < 0.001; probe 2, t(39) = 6.15, p < 0.001; probe 3, t(39) = 4.90, p < 0.001; probe 4, t(39) = 3.95, p < 0.001; probe 5, t(39) = 5.18, p < 0.001, **Figure 3G**; direct comparisons on the probes preceding the common problems, two-way repeated measures ANOVA, group X probe interaction, F(1, 78) = 78.62, p < 0.001; at probe 0, t(78) = 1.14, p = 0.26; at probe 5, t(78) = 8.61, p < 0.001; **Figure 3H**). These findings indicate that the model expressed during probe tests contained information that could be used to predict the solutions to discrimination problems that had not yet been encountered (i.e., a predictive model).

If mice retrieve a predictive model during learning, then their learning might be affected by the accuracy of the predictions. For example, an accurate model may reduce the amount of new information that needs to be acquired and incorporated into memory, while inaccurate predictions might interfere with performance. To test this, we asked how discrimination performance on the first day of training was related to the accuracy of the prediction measured during the preceding probe test. We found that accurate predictions were associated with greater preference for the rich tone on the first day of training (r = 0.68, p < 0.001; LME model with a fixed effect for prediction accuracy and a random effect for subject, t(478) = 19.58, p < 0.001; **Figure 3I**; last day, r = 0.22, p < 0.001, LME model, t(478) = 5.00, p < 0.001; **Figure 3J**) and faster learning (i.e. fewer days to criterion, r = −0.46, p < 0.001, LME, t(478) = 11.22, p < 0.001; **Figure 3K**), indicating that mice learned new problems more easily when they possessed a predictive model. Relatedly, accurate predictions were associated with *less* learning overall. Mice that retrieved accurate predictions showed a smaller change in their preference for the rich tone between the first and last day of training (r = −0.34, p < 0.001; LME, t(478) = 7.61, p < 0.001; **Figure 3L**), as well as a smaller change in their memory, as indicated by higher correlations between their responses on the probe tests before and after training (r = 0.54, p < 0.001; LME, t(478) = 14.19, p < 0.001; **Figure 3M**). Together, these findings indicate that predictive models improve learning by reducing the amount of new information that must be learned.

### Structured training reactivates CA1 ensembles active during prior learning

Our behavioral data indicate that mice retrieve memories of prior problems to support predictions and new learning. The hippocampus plays a crucial role in memory retrieval by reactivating ensembles (e.g., engrams) that were active during prior experiences (Liu et al., 2012; Goshen et al., 2011; Tanaka et al., 2014). To examine how ensemble reactivation in the hippocampus might support the formation of a predictive model in this task, we implanted GRIN lenses above the CA1 layer of the hippocampus in 13 Thy1-GCaMP6f mice, and imaged 4349 unique neurons with custom miniature microscopes (Jacob et al., 2018; **Figure 4A, B**) while mice performed the discrimination and foraging test described above (n = 8 Structured training mice, n = 5 Unstructured training mice; **Supplemental figure 2**). Imaging data was collected during the seven probe tests and the first and last day of each of the six training problems (19 sessions). To measure ensemble reactivation in our task, we registered active neurons across sessions (*CellReg*, Sheintuch et al., 2017; **Figure 4C; Supplemental figure 3**) and then quantified excess population overlap for every pair of sessions, defined as the number of neurons that were active in both sessions divided by the number that were active in either session (**Figure 4D**) minus the overlap expected due to the amount of time between imaging sessions (**Supplemental figure 4**). We found that excess overlap increased with Structured training, but not Unstructured training (LME model with fixed effects for sessions and group, and a random effect for subject, session X session X group interaction, t(4438) = 3.78, p < 0.001; post-hoc LME models with fixed effects for sessions, and a random effect for subject, Structured training group session X session interaction, t(2732) = 6.09, p < 0.001, Unstructured training group session X session interaction, t(1706) = 0.23, p = 0.81; **Figure 4E; Supplemental figure 4D, E**), indicating that Structured training increased the degree to which hippocampal ensembles were reactivated during later sessions. Interestingly, ensemble reactivation increased in the Structured training group at approximately the same time that these mice began to form a predictive model, suggesting that neuronal reactivation may be a mechanism for integrating experiences into a model (**Figure 2C-D**).

**Figure 4.**
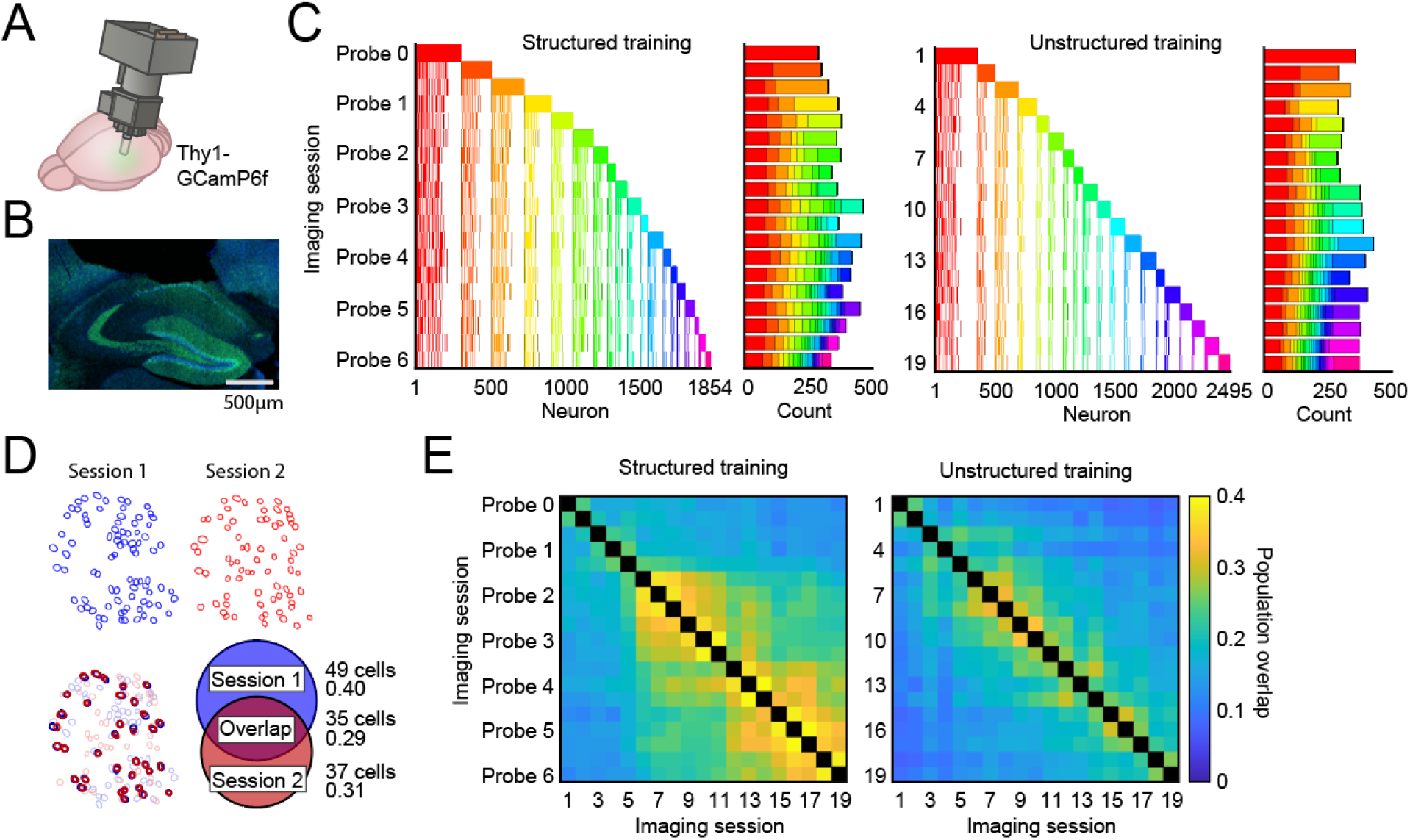
Mice reactivate CA1 populations during structured, but not unstructured, training. **(A)** Cartoon of the custom miniature microscope implanted into the hippocampus. **(B)** Histological section used to identify the location of a GRIN lens implanted on the surface of CA1. **(C)** We tracked the activity of every neuron throughout training for both the Structured training and Unstructured training groups (see also **Supplemental figure 3**). The columns show the activity of one neuron over all 19 training sessions (7 probes and the first and last day of each of 6 problems). Neurons are colored according to the session in which they were first active. Rows show the composition of every session in terms of which cells were active. The composition is summarized in the outset bar chart. **(D)** An explanation and example of the population overlap computation, defined as the number of cells active in both sessions divided by the number active in either. **(E)** The mean observed overlap between every pair of imaging sessions for the Structured training and Unstructured training groups. Matrices showing *excess* overlap are shown in **Supplemental figures 4D-E**.

### CA1 ensemble activity patterns stabilize during Structured training

In addition to activating unique ensembles of neurons, the hippocampus also guides memory retrieval by reorganizing neuronal activity patterns along dimensions that are important for task performance (i.e., remapping; Muller and Kubie, 1987; Markus et al., 1995; Smith and Mizumori, 2006; Kelemen and Fenton, 2010; Bulkin et al., 2016). To assess how the reorganization of hippocampal activity might support the formation of a predictive model in our task, we investigated time-warped neuronal activity during events common to every trial (from nose poke until the mouse waited in the food hopper for 2 seconds after tone-on; **Figure 5A**). Individual neurons reliably became active at discrete periods during every trial (**Figure 5B**) such that the population of neurons encoded the entire trial progression (**Figure 5C**). To evaluate how patterns of activity changed over training, we identified neurons that were active in multiple sessions and measured the similarity of their activity patterns between sessions (**Figure 5D**) while controlling for the effect of time between sessions as above (i.e., excess activity similarity, **Supplemental figure 4**). Comparing the excess activity similarity between every pair of sessions for both groups showed that population activity became more similar with Structured training only (LME model with fixed effects for sessions and group, and a random effect for subject, session X session X group interaction, t(3730) = 5.77, p < 0.001; post-hoc LME models with fixed effects for sessions and a random effect for subject, Structured training group session X session interaction, t(2334) = 7.23, p < 0.001, Unstructured training group session X session interaction, t(1396) = 1.57, p = 0.12; **Figure 5E; Supplemental figure 4**). Therefore, Structured training increased the degree to which hippocampal neurons from early sessions maintained their activity patterns when reactivated in later sessions, consistent with the idea that this activity supported the retrieval of training memories late in learning. As with the increase in ensemble overlap described above, this increase in activity similarity occurred at approximately the same time that the Structured training group formed a predictive model (**Figure 2C-D**). Together, this suggests that the hippocampus supported a predictive model by repeatedly reactivating prior training memories that predicted the answers to future problems and enabled efficient learning.

**Figure 5.**
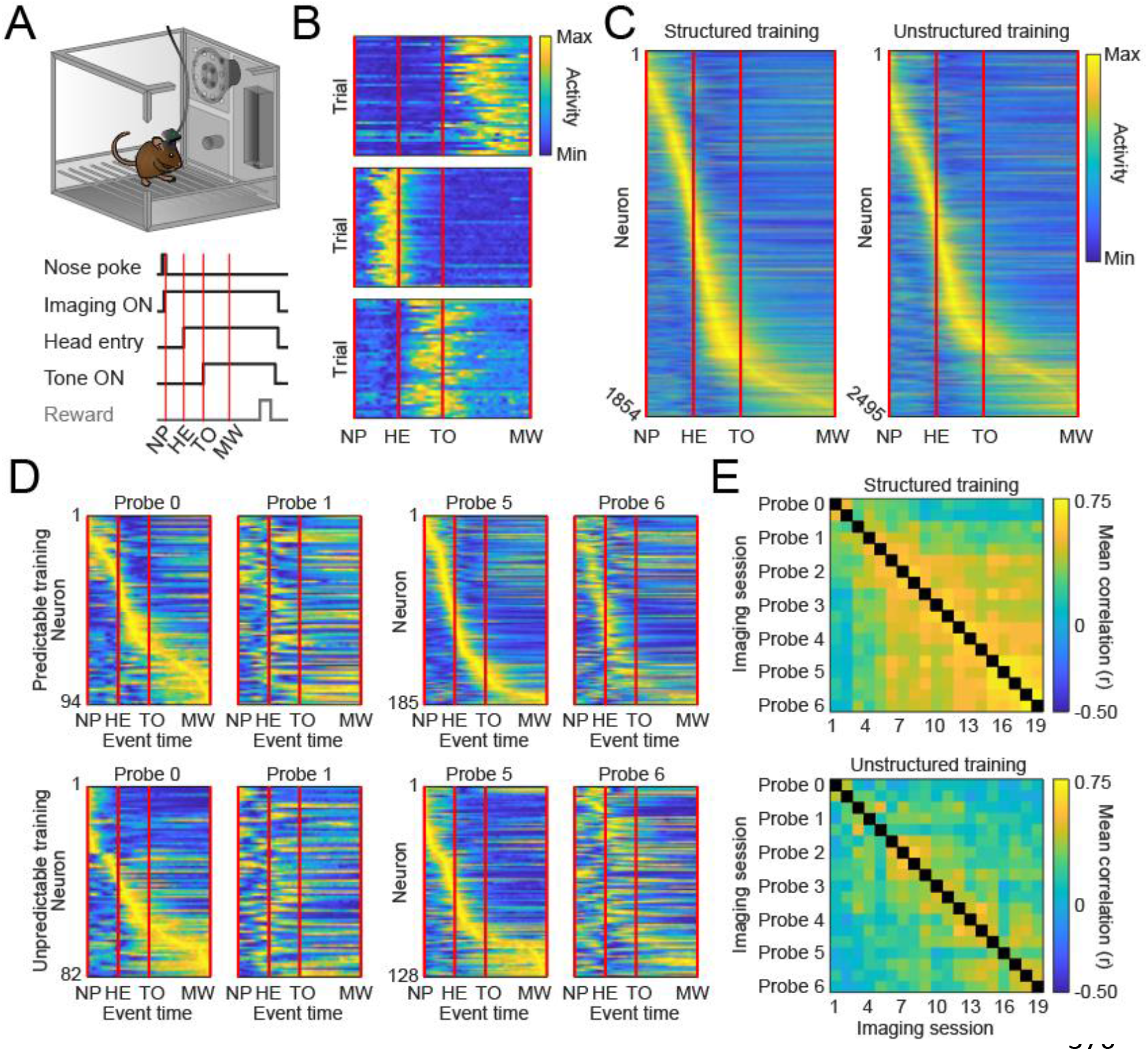
Mice reactivate CA1 activity patterns during Structured training. **(A)** We examined activity during the window between nose-poke and then end of the minimum wait (MW) period, occurring 2s after tone-on. **(B)** Three examples of neurons that were reliably active during discrete periods on every trial. **(C)** The mean activity is shown for every neuron imaged from each group, arranged by when during the trial it was most active. **(D)** The activity patterns of neurons that were active across any pair of probe sessions surrounding common problems (probes 0 & 1, 5 & 6) are shown. We examined the similarity of population activity across sessions by measuring the correlation between the activity patterns of individual cells in both sessions. The top row shows all neurons imaged from mice in the Structured training group that were active in both probe 0 and 1 (left side), and then neurons that were active in both probe 5 and 6 (right side). The bottom row shows neurons imaged from mice in the Unstructured training group. For each pair of probes, the neurons are sorted according to their activity in the left-side probe (probe 0 or probe 5), such that corresponding rows show the activity of the same neuron. Note the highly similar activity patterns between probes 5 and 6 in the Structured training group only. **(E)** The mean observed activity pattern similarity between every pair of imaging sessions for the Structured training and Unstructured training groups. Matrices showing excess activity similarity are shown in **Supplemental figure 4F-G**.

### Learning-related neuronal activity is incorporated into a task memory ensemble

Because the hippocampal ensemble stabilized at the same time that mice began to form a predictive model, we hypothesized that this stability reflected the transition from learning new problems to predicting them. In particular, we hypothesized that mice reactivated and updated ensembles during learning, and that the ensembles stabilized when mice developed a predictive model. To test this, we quantified the degree to which the ensembles from earlier sessions were reactivated during later sessions in terms of a reactivation score that described both the ensemble overlap between a pair of sessions and the similarity of their activity patterns (see methods; time-corrected results reported in **Supplemental figure 5**). We then tested the relationship between ensemble reactivation and behavior.

We first investigated whether mice reactivated prior ensembles and associated memories during new learning by comparing the ensemble reactivation scores between probe sessions and the training sessions one day later. Consistent with the above findings, we observed that the Structured training group increased ensemble reactivation during learning specifically during later training problems (Structured training, F(5, 35) = 6.35, p < 0.001), while the Unstructured training group did not (Unstructured training, F(5, 20) = 1.67, p = 0.19; **Figure 6A**; between groups comparisons, LME model with fixed effects for session and group, and a random effect for subject, session X group interaction, t(22) = 2.44, p < 0.05; **Figure 6B**). However, if ensemble reactivation is indicative of predictive memory retrieval, then the mice should show greater reactivation specifically when they show behavioral evidence of accurate memory retrieval. Consistent with this, we observed that ensemble reactivation scores were correlated with accurate discrimination performance on the first day of training, our behavioral measure of memory retrieval during learning (r = 0.39, p < 0.001; LME model with fixed effect for first day discrimination and a random effect for subject, t(75) = 3.52, p < 0.001; **Figure 6C**).

**Figure 6.**
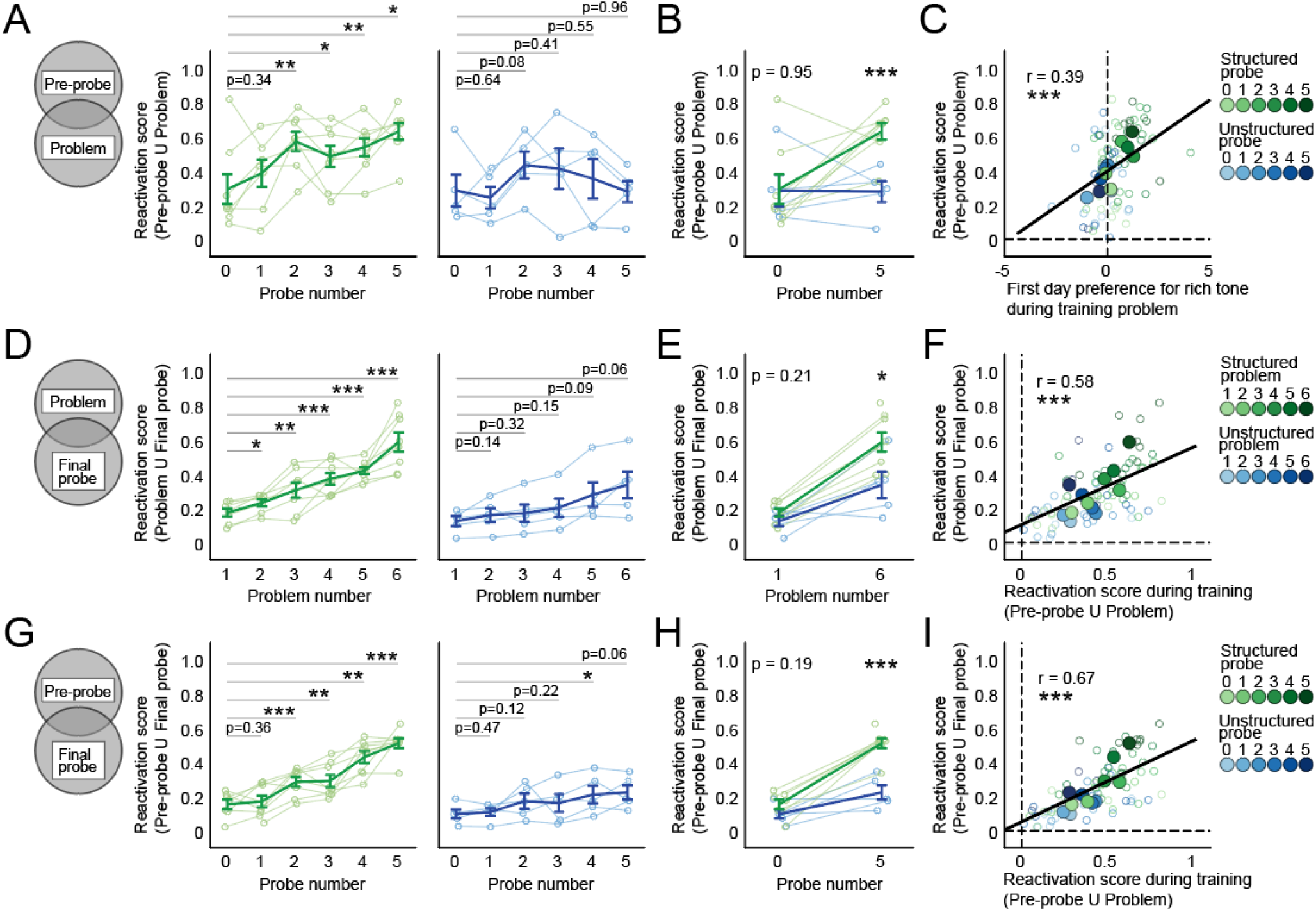
Learning-related neuronal activity is incorporated into a reactivated ensemble. **(A)** We computed a reactivation score that combined ensemble overlap and activity similarity measures. The reactivation probe session population during subsequent training problems is shown for the Structured training group (left) and the Unstructured training group (right). **(B)** The reactivation of probe ensembles during subsequent common problems is shown for both groups. **(C)** The reactivation of probe ensembles during subsequent problems was predicted by first day discrimination accuracy, a behavioral measure of predictive memory retrieval. Large circles show condition means. **(D)** The reactivation of training session ensembles during the final probe session is shown for the Structured training group (left) and the Unstructured training group (right). **(E)** The reactivation of the common training problem ensembles during the final probe session is shown for both groups. **(F)** The reactivation of training ensembles during the final probe was predicted by the reactivation of probe ensembles during training. **(G)** Like D, but with the reactivation of probe ensembles during the final probe session. **(H)** Like E, but with probe ensembles. **(I)** The reactivation of probe ensembles during the final probe was predicted by the reactivation of probe ensembles during training problems.

In addition to supporting memory retrieval, reactivating prior ensembles during learning may also support the formation of predictive memories by promoting memory updating and integration (Zeithamova et al., 2012; Cai et al., 2016; Rashid et al., 2016). To investigate the degree to which reactivated ensembles were updated with new neurons and activity patterns during training sessions, we compared the ensemble observed during the final probe test—when mice expressed their fully integrated predictive model—to the ensembles observed during individual training sessions. If training-session information is incorporated into the final predictive model, then there should be increased reactivation of the training ensemble during the final probe. Consistent with the evidence for increased reactivation in the Structured training group described above, we observed that the later Structured training sessions were most strongly reactivated during the final probe session (Structured training group, F(5, 35) = 27.18, p < 0.001; Unstructured training group, F(5, 20) = 5.64, p < 0.01; **Figure 6D;** between-group comparison, LME model with fixed effects for session and group, and a random effect for subject, session X group interaction, t(21) = 2.30, p < 0.05; **Figure 6E**). Analyses controlling for time between imaging sessions confirmed that only Structured training sessions showed excess reactivation scores that exceeded chance (i.e., what was expected due to the amount of time between imaging sessions), beginning with the second training session (**Supplemental figure 5G-H**). This indicates that Structured training promoted the development of a final probe ensemble that was comprised of neurons and activity patterns seen during earlier training problems.

Importantly, if the incorporation of training-related information into the predictive model is due to a memory updating process occurring during training, then the degree to which training ensembles are reactivated during the final probe should be related to neural processes occurring during training.

Specifically, we would expect to see greater updating during training sessions with stronger reactivation of the to-be-updated ensemble. Consistent with this, we found that reactivation during the final probe was correlated with the amount of reactivation observed during training: the greater the reactivation of the preceding probe ensemble during training, the greater the reactivation of that training ensemble during the final probe test (r = 0.58; p < 0.001; LME model with fixed effect for first day discrimination and a random effect for subject, t(74) = 5.52, p < 0.001; **Figure 6F**, **Supplemental figure 5I**). This suggests that the ensemble observed during the final probe was repeatedly reactivated during—and updated with new neuronal activity patterns from—each training problem.

If so, then the ensembles observed during each probe test should become progressively more similar to the final probe test ensemble as mice accumulate information from each training problem into a predictive model. Consistent with this, probe reactivation scores increased over training for both groups (Structured training group, F(5, 35) = 27.62, p < 0.001; Unstructured training group, F(5, 20) = 3.05, p < 0.05; **Figure 6G**), but this increase was greater in the Structured training group (LME model with fixed effects for session and group, and a random effect for subject, session X group interaction, t(21) = 4.21, p < 0.001; **Figure 6H**), and only the Structured training group showed excess reactivation over that expected by chance (**Supplemental figure 5J-K**). Similar to above, we also found a strong correlation between probe ensemble reactivation during training and probe ensemble reactivation during the final probe (r = 0.67, p < 0.001; linear mixed effects model with fixed effect for first day discrimination and a random effect for subject, t(74) = 7.51, p < 0.001; **Figure 6I**, **Supplemental figure 5L**), consistent with the idea that the incorporation of new information required ensemble reactivation during learning. Together, these data suggest that the increased ensemble stability observed in the Structured training group may be attributed to the repeated reactivation of a predictive model and corresponding ensemble, both formed by incorporating information from each of the training problems.

## Discussion

Hippocampal activity patterns are hypothesized to reflect predictive models of the environment (Sanders et al., 2020). Here, we tested this hypothesis by investigating the relationship between memory, prediction, and CA1 ensemble activity as mice learned the rules that governed their environment. By using a new behavioral method for observing how memories change after discrete learning experiences, we found that Structured learning experiences promoted the integration of new learning into a memory of the task environment that accurately predicted future problems (i.e., a predictive model). We showed that this model included predictions about untrained stimuli, and that the accuracy of these predictions had a large effect on the efficiency of new learning. Then, by observing hippocampal activity throughout learning, we found that mice learning Structured training problems reactivated prior ensembles, thereby supporting the integration of new information into the reactivated ensemble. These data indicate that hippocampal ensemble reactivation supports the formation of predictive models that explain the underlying structure of an environment. Moreover, these data shed light on how memory systems organize experiences to maximize prediction accuracy and minimize memory interference.

The hippocampus has long been implicated in memory processes that support predictive models (Tolman, 1948) and considerable evidence suggests that hippocampal activity develops as subjects learn about their environment (Smith and Mizumori, 2006; Gill et al., 2011; Plitt and Giocomo, 2021). However, a fundamental feature of a predictive model is the ability to infer information beyond what has been directly experienced—to generalize what has been learned to explain and predict aspects of the environment that have not been learned (Tervo et al., 2016; Whittington et al., 2018, 2022). Consistent with this, we found that mice readily generalized what they learned from individual discrimination problems into predictions about novel contingencies during subsequent probe tests. Predictions expressed during probe tests, in turn, affected how mice learned new problems, such that accurate predictions accelerated learning and inaccurate predictions impeded learning. These findings are consistent with longstanding ideas that it is easier to learn things that are related to what is already known (Bartlett, 1932; Bransford and Johnson, 1972; Craik and Tulving, 1975; Harlow, 1949; Chase and Simon, 1973; Tse et al., 2007). These findings also highlight the reciprocal relationship between learning and memory whereby learning is stored in memory and then retrieved during new learning.

We additionally found that the Structured training group successfully integrated new experiences into a predictive model, whereas the Unstructured training group did not. This indicates that the ability to integrate experiences was influenced by whether they shared an underlying structure. It has been previously noted that the relationship between new experiences and existing memories plays an important role in memory formation (Fernández et al., 2016; van Kesteren et al., 2012). However, it has not always been clear what makes a given experience sufficiently related to another to promote memory integration. One possibility is that mice selectively integrate information that is predicted by the memories retrieved during learning. Such a rule could support the formation of predictive models by organizing experiences in memory according to shared hidden causes (i.e. whether both experiences were caused by the same underlying rule or structure). It is also likely that memory integration in the Structured training group was facilitated by the learning-related benefits of retrieving accurate predictions: predicted experiences are learned more quickly, thereby increasing the likelihood that learning occurs in the presence of the retrieved memory. Inversely, if a memory interferes with learning (as was observed in both groups early in training and in the Unstructured training group throughout training), then the retrieved memory may be suppressed, thereby preventing the new information from being integrated into it. Over many iterations, these two processes could organize a series of experiences into a predictive model that closely matches the underlying structure of the environment and can therefore support predictions about future experiences in that environment.

Integrating experiences requires a system for retrieving relevant memories and then updating these memories with new information. The hippocampus plays essential roles in memory retrieval (Liu et al., 2012; Robinson et al., 2020) and consolidating new learning (Davis and Squire, 1984; Girardeau et al., 2009). One way these functions cooperate is through the process of integrating memories that are learned under the same hippocampal activity pattern, such as when two experiences occur close in time (Cai et al., 2016; Chowdhury et al., 2022) or when a second experience reminds one of the first (Zeithamova et al., 2012; Lee et al., 2017). In the current study, we found increased reactivation of hippocampal activity patterns associated with prior problems exclusively in the Structured training group and specifically late in learning, coinciding with the time that memory integration became apparent behaviorally. This suggests that memory integration occurs when predictive memories and their corresponding hippocampal activity states are retrieved during new learning, leading to integration of newly learned information into the reactivated state. If so, then the memory integration might result from a two-part process whereby memory retrieval increases the activity of the subset of neurons associated with the retrieved memory (i.e., the engram), and then a neuronal allocation process preferentially assigns newly learned information to those active neurons (Rashid et al., 2016). This process would result in the first and second experience becoming linked in memory due to both experiences sharing a single neuronal population that drives the retrieval of both memories when reactivated.

A complementary possibility is that the hippocampus changes its activity state in response to unpredicted problems. Although retrieving related memories is important for memory integration and the formation of predictive memories, it is arguably even more important to avoid retrieving and integrating unrelated information, which can cause memory interference (Smith and Vela, 2001) and possibly amnesia (McCloskey and Cohen, 1989). The hippocampus is well known for its ability to reorganize its firing activity in response to new environments (Muller and Kubie, 1987). Interestingly, this type of reorganization can also free subjects from memory interference (Bulkin et al., 2016; Butterly et al., 2012), likely by discouraging the retrieval of the interfering information (Park et al., 2021). Therefore, one explanation for our finding that the Unstructured training group showed little reactivation of prior hippocampal activity patterns during learning is that the hippocampus actively generated new patterns to facilitate the learning of unstructured problems. This would also help to avoid erroneously integrating experiences that do not share an underlying hidden cause and whose integrated memory would therefore have little predictive value.

Memory organization can also occur ‘offline’ during sleep or otherwise outside of task performance. In the days and weeks after learning, newly learned information goes through a systems consolidation process that transfers information into neocortical regions (Squire, 2004), and there are changes to memory content such as the loss of detailed information (Wiltgen and Silva, 2007; Richards et al., 2014) and, in some cases, additional learning or insight (Barron et al., 2020; Payne et al., 2009). Mice in the current study learned training problems over the course of weeks (**Figure 1F-G**), providing sufficient opportunity for long-term memory generalization into the cortex (Makino and Komiyama, 2015; Miller et al., 2019; Takehara-Nishiuchi and McNaughton, 2008; Tse et al., 2007, 2011). However, our data show that the process also occurs locally in the hippocampus. One possibility is that hippocampus supports the selective retrieval of predictive memories stored in the neocortex in order to update them with relevant new information (Debiec et al., 2002).

Together, these data suggest a concise description of prediction-based memory organization in the hippocampal system. After any learning experience, the resulting memory trace generalizes to form predictions about novel stimuli that can be retrieved when those stimuli are encountered in the future. When the generalized memories are retrieved during new learning, they are accompanied by the reactivation of hippocampal activity patterns that were present when the original memory was learned. If the retrieved memory improves learning, then the prior hippocampal activity patterns remain activated while the new memory is formed, thereby integrating the two memories. Alternatively, if the retrieved memory impedes learning, then the hippocampus changes its activity pattern to discourage the retrieval of the interfering memories and to avoid erroneous integration. Over many learning experiences, a predictive model is formed that matches the statistics of the experiences from which it was formed, and that can be used to predict other similar future experiences. Memories that are not incorporated into the model may be used to seed new models, or they may become isolated and forgotten.

## Acknowledgements

A.M.P.M. and P.W.F. designed the experiments and wrote the first draft of the paper. A.M.P.M., A.I.R. and M.L.D. performed the surgeries and did the histology. A.M.P.M. and A.D.J. designed and updated the behavioral and imaging hardware. A.M.P.M. conducted the experiments and performed the analyses. A.M.P.M., P.W.F. and S.A.J. supervised the project. All authors helped edit the paper. We thank Tao Zhang, Shuo Huang, Sandy Ma, and Lulu Liu for assistance with pilot versions of this project, Chen Yan, Lina Tran, and Andrew Mocle for assistance adapting the microscopes and imaging analysis pipeline, and Hendrick Steenland for designing components of the imaging acquisition hardware. We thank Nathan Insel for helpful comments on the manuscript. This work was supported by a Brain Canada platform grant and NIMH grant (RO1MH119421) to S.A.J. and P.W.F. A.M.P.M. was supported by a Restracomp award and a Research Institute Exceptional Trainee Award Fund Bursary from The Hospital for Sick Children. A.D.J. was supported by an Ontario Graduate Scholarship. A.I.R was supported by an NSERC CGS-D award and an NIH 1 F31 MH120920-01 award. M.L.D. was supported by a CIHR Vanier Canada Graduate Scholarship.

---

**Supplemental figure 1.**
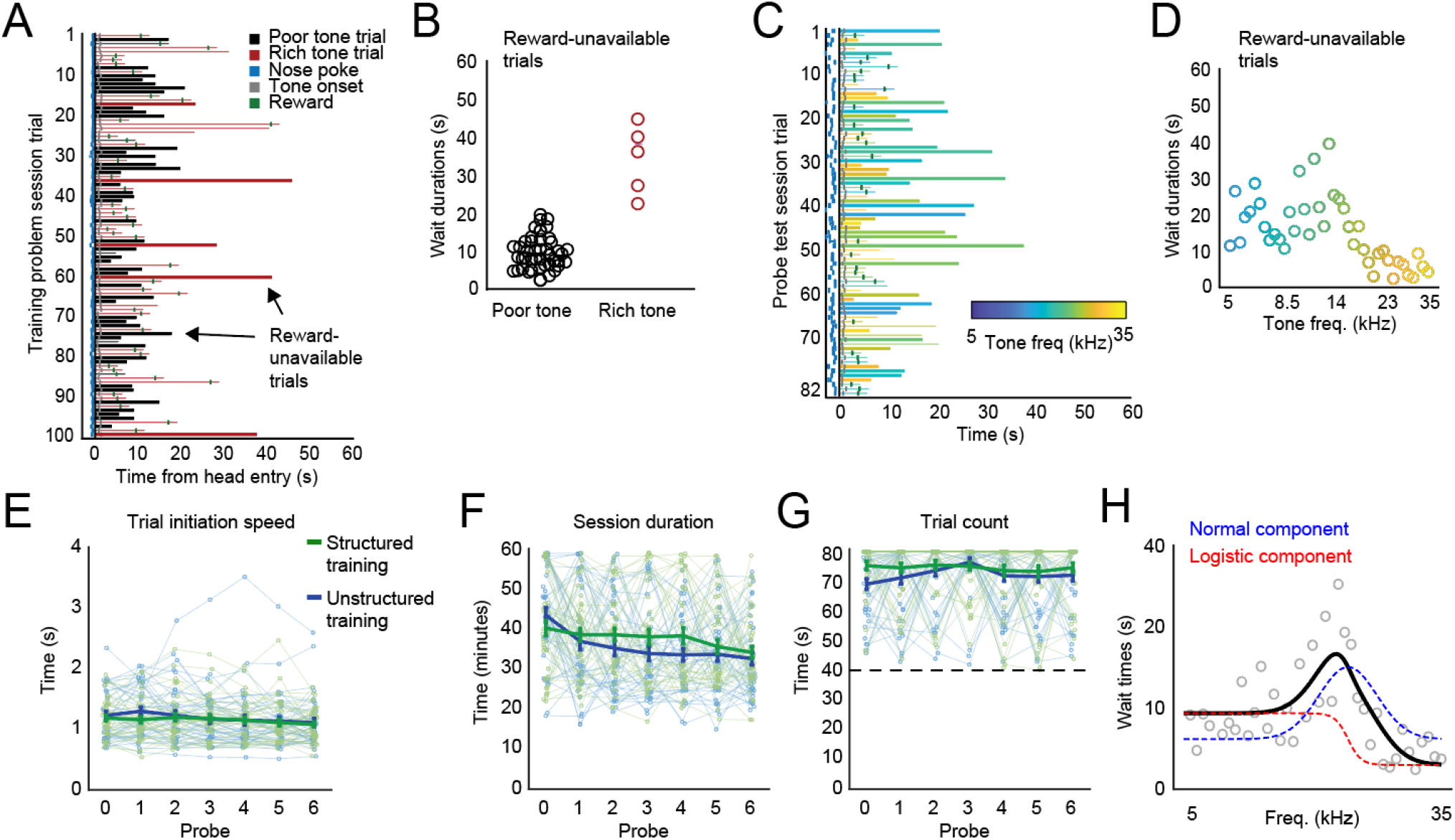
Operant behavior in unoperated mice. **(A)** Every measured behavior during an exemplary training session. Mice had the potential to earn 50 rewards over 100 trials in an hour-long session. On each trial, one of two unique tones would play: the rich tone (p(reward)=0.9) or the poor tone (p(reward) = 0.1) and the mouse responded by waiting for a potential reward. The wait durations on trials when no reward was available (thick lines) indicated the mouse’s preference for the tone that played on that trial. **(B)** All wait durations from reward-unavailable trials in A. **(C)** Every measured behavior during an exemplary probe test session. Mice had the potential to earn 41 rewards over 82 trials in an hour-long session. On each trial, one of 41 unique tones would play. No reward was available on half of all trials (all tones p(reward) = 0.5) and the wait durations on these trials (thick lines) indicated the mouse’s preference for each tone. In this example of behavior on probe 1 (after training on problem 1), the mouse waits longer in response to lower-frequency tones **(D)** All wait durations from reward-unavailable trials in C. **(E)** The speed of task behavior was similar between the groups throughout learning. The amount of time between nose poke entry and head entry is shown for every subject at every probe session. **(F)** The amount of time required to complete a session is shown for every probe session. Probe sessions were completed after 82 trials with a maximum time of 60 minutes. **(G)** The number of trials completed in every probe session is shown. Sessions with fewer than 41 trials (dotted line) were repeated. **(H)** For some analyses (**Figure 3; Supplementary figure 2**), we estimated the preference for every tone by fitting a curve composed of a normal component and a logistic component to the observed wait times on every trial (probe sessions only; see methods).

**Supplemental figure 2.**
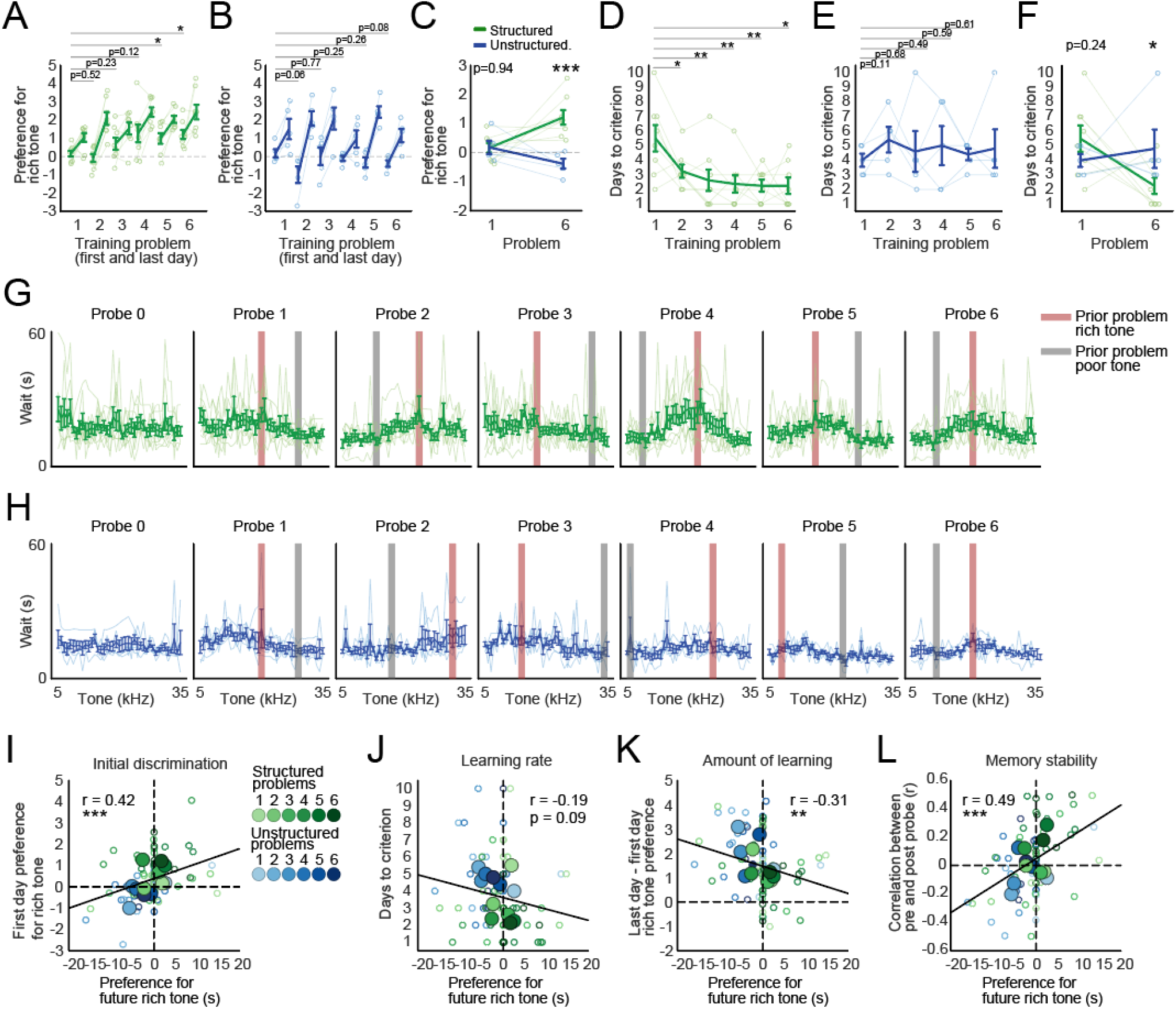
Operant behavior during calcium imaging. **(A)** Thirteen mice (8 Structured training, 5 Unstructured training) were imaged performing the discrimination and foraging task in an operant chamber. Preference for the rich tone (standardized mean difference between wait times in response to rich and poor tones) is shown for the first and last day of training on every training problem. The Structured training group showed a greater preference for the rich tone on the first day of training on problems later in learning (one-way repeated measures ANOVA, F(5,35) = 3.35, p < 0.05). **(B)** The Unstructured training group never improved its first-day preference for the rich tone (one-way repeated measures ANOVA, F(5,20) = 1.59, p = 0.21). **(C)** Direct comparisons between the two groups on the first and last problem revealed that the structured group showed greater preference for the rich tone on the first day of the last problem (mixed model with fixed effects for intercept, training group, problem number, and group X problem and a random effect for subject, group X problem coefficient, t(22) = 3.97, p < 0.001). **(D)** The number of days required to reach criterion is shown. The Structured training group learned new problems faster later in learning (one-way repeated measures ANOVA, F(5,35) = 6.16, p < 0.001). **(E)** The Unstructured training group never improved its learning rate (one-way repeated measures ANOVA, F(5,20) = 0.28, p = 0.92). **(F)** Direct comparisons between the two groups on days required to reach criterion on the common problems (group X problem coefficient, t(22) = 2.64, p < 0.05). **(G)** The wait time responses to every tone are shown for every subject in every probe test of the Structured training group. **(H)** The same as G, but for the Unstructured training group. **(I)** We measured predictions from behavior during the probes preceding each problem (see **Figure 3E**). The more accurate the prediction, the better the discrimination performance on the first day of training (linear effects mixed model with fixed effects for intercept, prediction accuracy, and a random effect for subject: prediction accuracy term, t(76) = 3.33, p < 0.01). Large dots indicate condition means. **(J)** The relationship between prediction and learning rate on the training problem (prediction accuracy term, t(76) = 1.92, p = 0.06). **(K)** The more accurate the prediction, the less new information was learned during training (prediction accuracy term, t(76) = 2.35, p < 0.05). **(L)** The more accurate the prediction, the more stable the memory as measured during pre and post probe tests (prediction accuracy term, t(76) = 4.53, p < 0.001).

**Supplemental figure 3.**
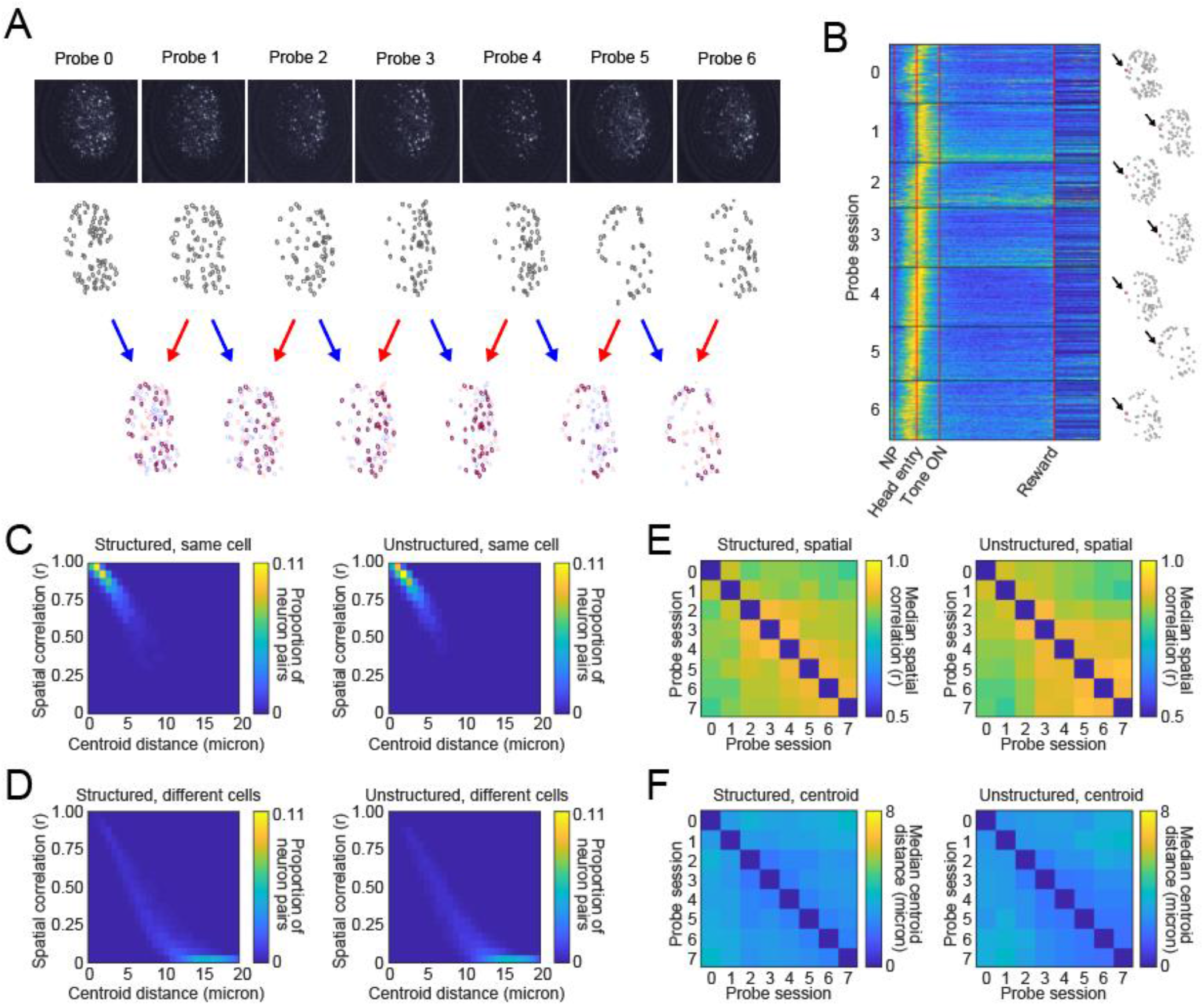
Registering CA1 neurons. **(A)** Example showing the registration of imaging data collected over weeks. Top row, fields of view (maximal projection images) from the same mouse from every probe session. Second row, spatial footprints are identified from the video file and aligned (translated and rotated) to a common reference frame. Bottom row, session pairs are then overlaid to identify overlapping neurons. **(B)** Example of a neuron that was active during every probe 0 – 6 and showed a stable activity pattern. Rows show normalized activity on every trial from every probe session. The neuron’s spatial footprint is highlighted to the right of each probe session. **(D)** Anatomical distance vs spatial footprint similarity for every registered (same cell) cell pair in the dataset. For each cell pair, we plotted the distance between the two cells versus the correlation coefficient of their cell mask spatial correlations. **(E)** Same as B, but for unregistered (different cells) cell pair. **(E)** Median spatial correlations for every registered cell pair from each probe session pair for each group. **(F)** Same as E, but with anatomical distance between centers of each cell.

**Supplemental figure 4.**
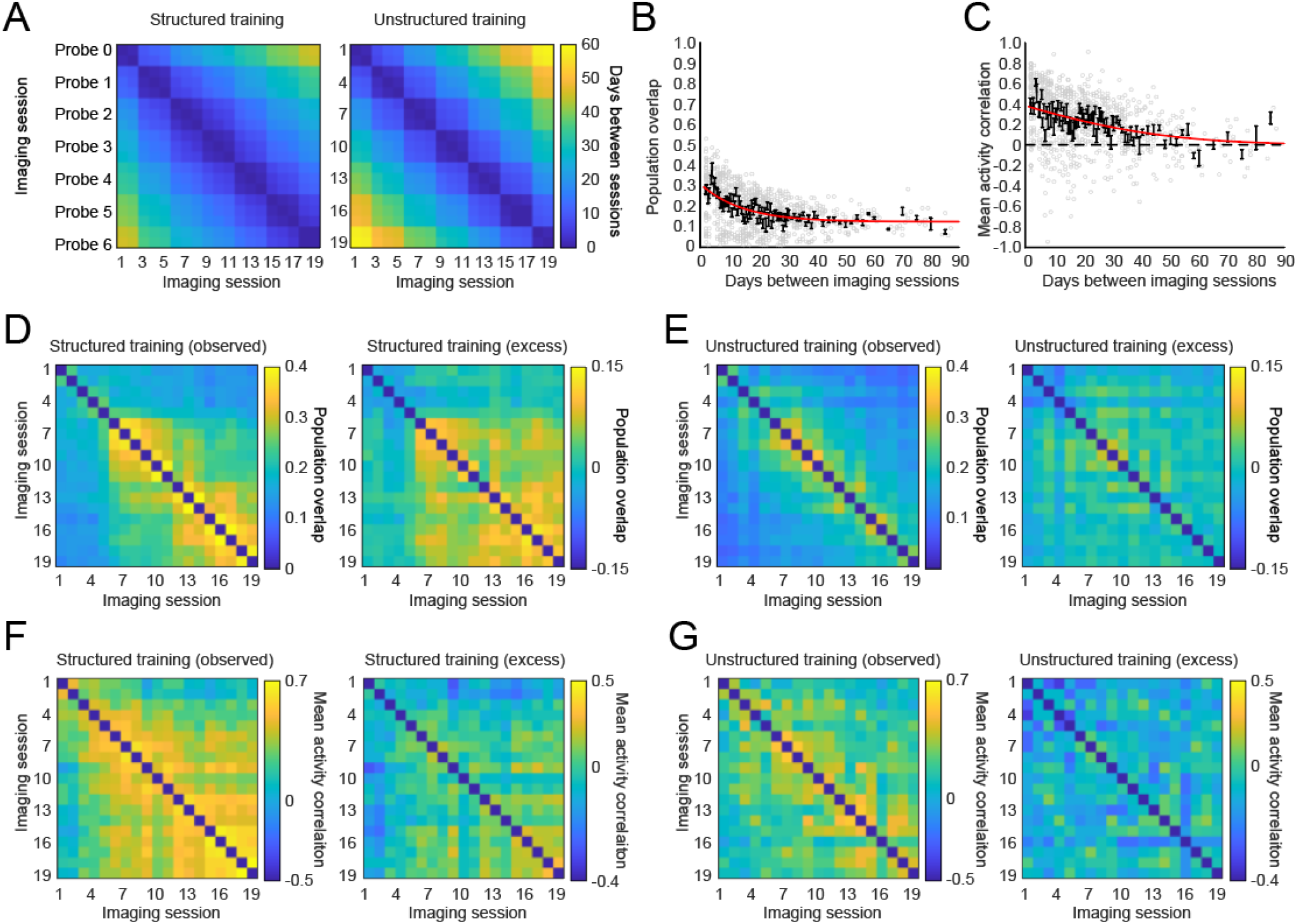
The relationship between ensemble similarity and time. **(A)** The average time between every imaging session is shown for both the structured and Unstructured training groups. **(B)** We modelled the effect of time on reactivation scores by fitting a truncated normal distribution to the ensemble overlap and mean activity similarity values obtained by comparing all pairwise combinations of imaging sessions from the Unstructured training group. Mean overlap in bins spanning time is shown in black. The fit model curve is shown in red. We obtained the time-corrected (excess) overlap values by subtracting the effect characterized by the fit curve from the observed overlap values. **(C)** Like B, but with the mean activity similarity across cells that were active any two sessions. **(D)** The ensemble overlap between every pair of sessions during Structured training is shown in terms of the observed values and the excess values. **(E)** Same as C, but for Unstructured training. **(F-G)** Like D and E, but with mean activity similarity.

**Supplemental figure 5.**
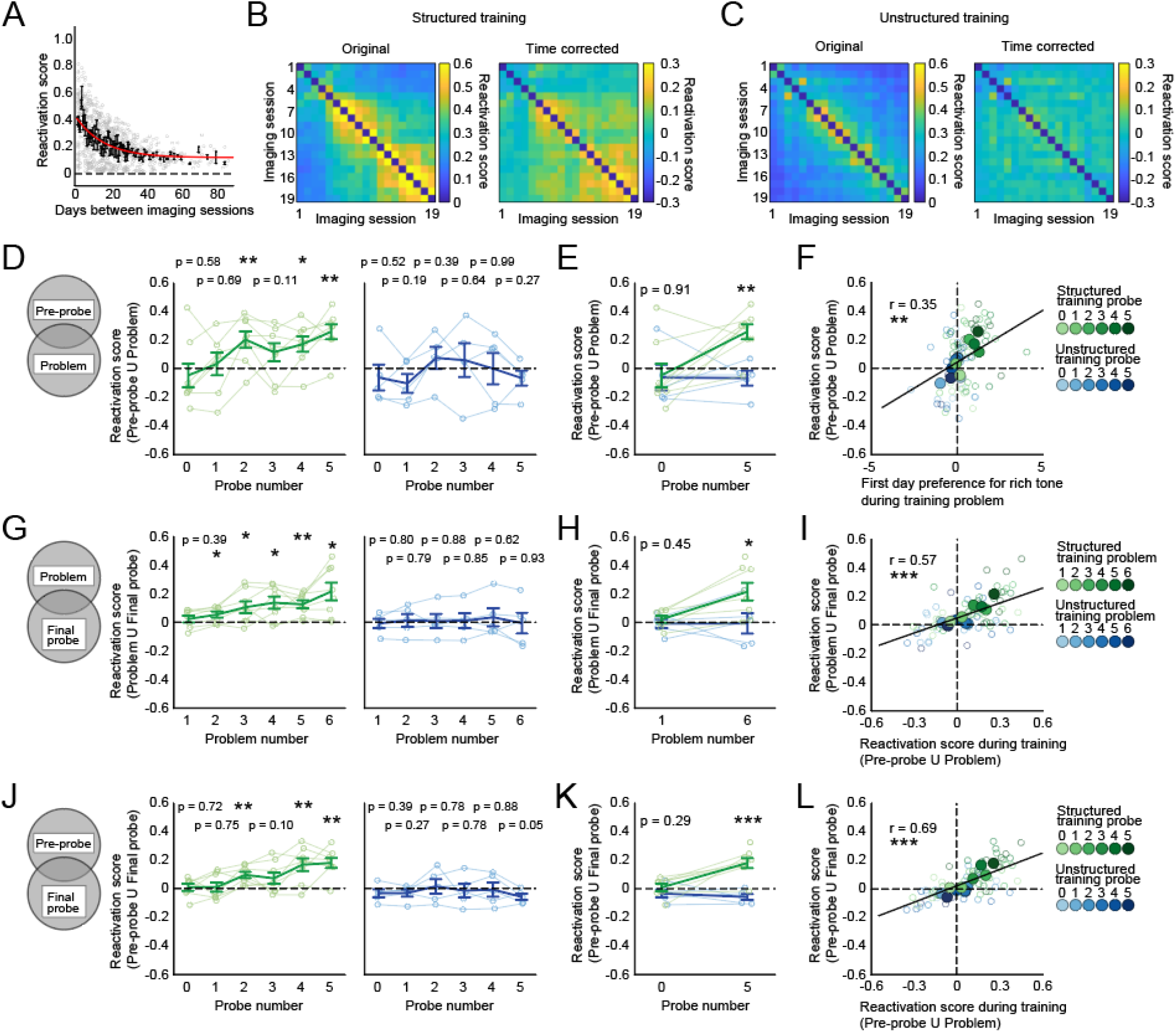
Learning related neuronal activity is incorporated into a reactivated ensemble, time corrected. **(A)** We modelled the effect of time on reactivation scores by fitting a truncated normal distribution to the reactivation scores obtained by comparing all pairwise combinations of imaging sessions from the Unstructured training group. **(B)** The observed and time-corrected mean reactivation score values for all pairwise Structured training group imaging session comparisons. **(C)** The observed and time-corrected mean reactivation score values for all pairwise Structured training group imaging session comparisons. There was a significant group (structured vs Unstructured training) X session 1 X session 2 interaction for the time-corrected reactivation scores (linear mixed model with fixed effects for session 1, session 2, and group, and a random effect for subject, t(4344) = 7.34, p < 0.001). **(D)** The reactivation probe ensembles during subsequent training problems increased with training for the Structured training group (F(5,35) = 5.39, p < 0.001; left) and but not the Unstructured training group (F(5, 20) = 1.55, p = 0.22; right). **(E)** The reactivation of probe ensembles during subsequent common problems are shown for both groups. Only the structured group showed an increased reactivation score late in learning (fixed effects for group and session and a random effect for subject, group X session interaction, t(22) = 2.37, p < 0.05). **(F)** The reactivation of probe ensembles during subsequent problems was predicted by first day discrimination accuracy (fixed effect for discrimination accuracy and a random effect for subject, t(75) = 3.15, p < 0.01). **(G)** The Structured training group showed increased reactivation of later training session ensembles during the final probe session (F(5,35) = 6.12, p < 0.001; left), while the Unstructured training group did not (F(5,20) = 0.24, p = 0.93, right). **(H)** Only the Structured training group showed an increase at the common training problems (fixed effects for group and session and a random effect for subject, group X session interaction, t(21) = 2.18, p < 0.05). **(I)** The reactivation of training ensembles during the final probe was predicted by the reactivation of probe ensembles during training (fixed effect for training reactivation and a random effect for subject, t(21) = 2.18, p < 0.05). **(J)** Like G, but with the reactivation of probe ensembles during the final probe session (Structured training, F(5,35) = 6.94, p < 0.001; Unstructured training, F(5,20) = 0.92, p = 0.49). **(K)** Like H, but with probe ensembles (fixed effects for group and session, group X session interaction, t(21) = 3.75, p < 0.01). **(L)** The reactivation of probe ensembles during the final probe was predicted by the reactivation of probe ensembles during subsequent training problems (fixed effect for training reactivation and a random effect for subject, t(74) = 7.40, p < 0.001).

## Methods

### Animals

All experiments conformed to the guidelines set forth by the Canadian Council for Animal Care and the Animal Care Committee at the Hospital for Sick Children. For the behavior-only study, we used 80 wild type mice aged 8-10 weeks at the start of training. These mice were derived from crossing C57BL6n and 129SvEv mice (breeding mice were sourced directly from Taconic). For the imaging study, we used 13 transgenic mice expressing the fluorescence calcium indicator GCaMP6f under the Thy1 promoter (GP5.17 mice, Jackson Laboratories, stock no. 025393; Dana et al., 2014). All mice were weaned at 21d, housed 4 to a cage, and maintained on food restriction at 80-85% of their free-feeding weight for the duration of the experiment. The housing room was maintained on a 12h-12h light-dark cycle with the lights coming on at 8am. All experiments were performed between 9am and 6pm. Cages of mice were randomly assigned to experimental groups.

### GRIN lens implantation surgery for calcium imaging

Mice were pretreated with atropine sulfate (0.1mg/kg, ip), anesthetized with chloral hydrate (400mg/kg, ip), injected with dexamethasone (5mg/kg, ip), and placed in a stereotaxic frame. An incision was made to expose and clean the skull, and then a craniotomy was drilled above the right dorsal hippocampus (AP = −2.0mm, ML = 1.5mm from bregma). To gain access to CA1, the cortex and corpus callosum above the hippocampus were gently aspirated while the craniotomy was constantly irrigated with artificial cerebrospinal fluid. A baseplate with an attached 2.0mm GRIN objective lens was slowly lowered into the craniotomy to a depth of 1.5mm below the surface of the skull and secured with jeweler screws and dental cement. The mice were then treated subcutaneously with meloxicam (5mg/kg) for analgesia. Imaging and behavior began 3-6 weeks later when calcium activity was observed.

### Behavioral apparatus

Operant conditioning chambers (Med Associates, VT, USA, ENV-307W-CT) were enclosed in sound attenuating cubicles (ENV-016MD). A tall liquid delivery cup (Med Associates, ENV-303LPHDW-4.25) was recessed in the center of the front wall and a nose poke (Med Associates, ENV-313W) was placed to the left of the liquid delivery cup. Both the liquid delivery cup and the nose poke were outfitted with infrared beams to detect head entries. A tweeter speaker (Med Associate, ENV-324D; pure tone range from 5 - 35 kHz) was mounted above the nose poke. We 3D printed custom face panels to improve functionality for mice with head-mounted microscopes. During imaging, the ceiling of the operant chamber was removed, and a small hole was opened at the top of the sound attenuating chamber to run imaging cables between the microscope and a commutator (NeuroTek Inc., ON, Canada). Operant boxes were controlled with custom code (Med Associates, Med-PC) running on a desktop computer in an adjacent room.

### Auditory stimuli

Mice were exposed to 41 logarithmically spaced tone frequencies that spanned their auditory range. The frequencies for all subjects were (1) 5000, (2) 5249, (3) 5511, (4) 5786, (5) 6074, (6) 6377, (7) 6695, (8) 7028, (9) 7379, (10) 7747, (11) 8133, (12) 8538, (13) 8964, (14) 9411, (15) 9880, (16) 10372, (17) 10890, (18) 11432, (19) 12002, (20) 12601, (21) 13229, (22) 13888, (23) 14581, (24) 15307, (25) 16070, (26) 16872, (27) 17713, (28) 18596, (29) 19523, (30) 20496, (31) 21518, (32) 22590, (33) 23716, (34) 24899, (35) 26140, (36) 27443, (37) 28811, (38) 30247, (39) 31755, (40) 33338, (41) 35000 kHz. To avoid mice using volume information when learning to discriminate between tones, we applied a volume correction based on R-weighting (Björk et al., 2000) to account for how hearing sensitivity varies across frequencies. The resulting volumes for each tone in decibels were (1) 83, (2) 83, (3) 83, (4) 83, (5) 83, (6) 83, (7) 83, (8) 83, (9) 83, (10) 83, (11) 83, (12) 83, (13) 83, (14) 82, (15) 81, (16) 81, (17) 80, (18) 79, (19) 79, (20) 78, (21) 77, (22) 76, (23) 76, (24) 75, (25) 74, (26) 74, (27) 73, (28) 73, (29) 73, (30) 72, (31) 72, (32) 73, (33) 74, (34) 74, (35) 75, (36) 76, (37) 77, (38) 78, (39) 81, (40) 83, (41) 83. Tone volume was also randomly modified +/− 5% on each trial. All frequencies and volumes were controlled by a dedicated programmable audio generator (Med Associates, ANL-926).

### Behavioral procedures

#### Pretraining

Pretraining involved a series of training days that first shaped the mice to nose poke for 0.04 mL chocolate milk rewards delivered in the adjacent food cup, and then shaped them to wait longer and longer for rarer and rarer rewards. The first day of nose poke training involved placing hungry mice into the operant box, with chocolate milk smeared on the nose poke port and in the liquid delivery cup. Chocolate milk was then automatically delivered at 1 min +/− 30s intervals, while nose pokes were reinforced with chocolate milk delivery in the liquid delivery cup. On subsequent days, chocolate milk delivery was contingent on a nose poke followed by a head entry into the liquid delivery cup within 5s. We shaped the mice to steadily increase the amount of time they were willing to wait for a reward after head entry from 0s to approximately 12s over days by increasing the time between head entry and reward delivery. By the end of pretraining, wait times on every trial were randomly drawn from a distribution based on a negative power distribution (min = 0 s, max = 56.80 s, mean = 6.35 s, median = 2.69 s). At the same time that wait times were increasing, the probability of receiving a reward given a sufficient wait response was steadily decreased from 100% to 50%. Pretraining typically lasted between 11 and 20 days.

#### Session order and progression criteria

After pretraining, mice began the task protocol. Mice were trained on six training problems (problems 1-6) interleaved with 7 probe tests (probes 0-6). Training began with a pre-probe (probe 0) to test for any pre-existing tone preferences. This was followed by the first problem (problem 1) and then probe 1, followed by problem 2 and then probe 2, and so on until the last problem (problem 6), which was followed by the final probe (probe 6). Probe sessions were administered on a single day, whereas problem sessions were administered daily until the mice achieved a behavioral criterion (described below). After achieving criterion, mice were given one additional day of training (i.e., the last day of training), after which they progressed to the probe test.

#### Training problem sessions

During a training problem session, mice learned to wait longer in response to a rich tone than to a poor tone. A trial began when the mouse nose-poked and then entered their head into the food cup within 4s, waited until the tone started playing (1 +/− 0.5s after head entry) and then waited for an additional 2s (i.e., the minimum wait). After achieving the minimum wait, a mouse could abandon the trial by exiting the food cup or it could wait for a delay of 0-60s for a possible reward. However, successfully waiting for the duration of the delay did not ensure a reward, since rewards were only available on 50% of trials on average. Therefore, mice learned to use the tones to determine how long to wait (i.e., how much time to invest on that trial) for a potential reward. On each trial, the tone was randomly selected from two possible tones: the “rich” tone, which indicated that the trial had a 90% chance of having an available reward, and the “poor” tone, which indicated a 10% chance. The tone frequencies for every problem are shown in **Table 1** (also see **Figures 1B-C**). We measured the animal’s preference for a tone in terms of how long it was willing to wait on trials when that tone was playing but no reward was available. We defined successful discrimination learning (i.e., the behavioral criterion) as significantly longer waits in response to the rich tone than to the poor tone as indicated by a significant t-test with an alpha level of 0.01 (see **Supplemental figure 1A-B**). Individual sessions lasted for 100 trials or 1 hour, whichever came first. Sessions with fewer than 41 trials were repeated. The number of days required to learn a problem varied with the prior experience of the animal and can be seen in **Figure 1F**.

**Table 1:**
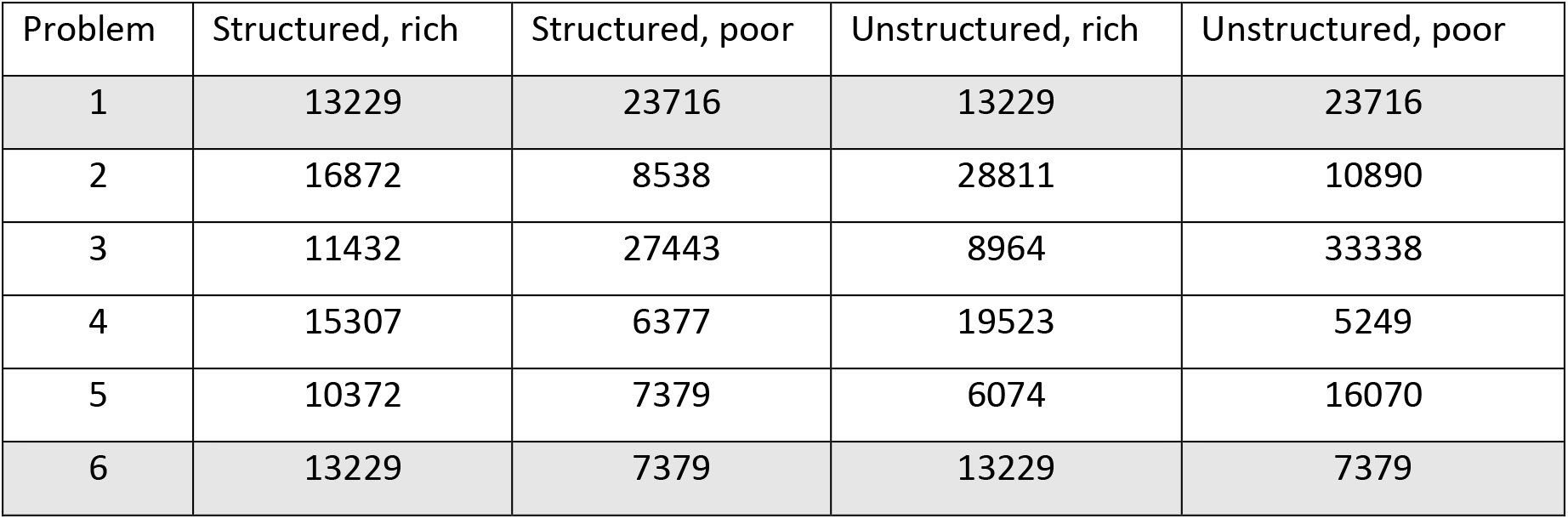
The tone frequencies used for each training problem. All frequencies are shown in kHz. The tones used in problem 1 were common between both groups, as were the tones used in problem 6 (both rows are shaded). To control for presentation order, half of the subjects were trained on the problems in reverse order.

| Problem | Structured, rich | Structured, poor | Unstructured, rich | Unstructured, poor |
| --- | --- | --- | --- | --- |
| 1 | 13229 | 23716 | 13229 | 23716 |
| 2 | 16872 | 8538 | 28811 | 10890 |
| 3 | 11432 | 27443 | 8964 | 33338 |
| 4 | 15307 | 6377 | 19523 | 5249 |
| 5 | 10372 | 7379 | 6074 | 16070 |
| 6 | 13229 | 7379 | 13229 | 7379 |

#### Probe test sessions

Probe test sessions were identical to training problem sessions except that during a probe test session, mice were exposed to all 41 tones used in the study to test their preferences. Tones were presented in random order, twice each, such that a reward was available on exactly one trial for each tone, resulting in an average reward probability of 50% (the same as in the problem training sessions). There was no behavioral criterion during probe test sessions. Individual sessions lasted for 82 trials or 1 hour, whichever came first. Each probe test (0-6) was only administered once unless the mice performed fewer than 41 trials, at which point the session was repeated.

### Calcium imaging

Imaging mice were acclimated to the weight of the imaging scope and cables during pretraining by slowly introducing heavier and heavier dummy scopes as the mice learned to nose poke for a reward. During the training protocol, we only collected video data during probe tests and during the first and last session of each training problem. To minimize the effect of bleaching over the course of a session, the LED light was kept off except for the period on each trial between nose poke and two seconds after reward delivery. If a mouse abandoned a trial before reward delivery, then the LED was turned off as soon as they left the food cup. The microscope LED was controlled by the MED-PC code so that the ON and OFF times were synchronized with task performance.

To encourage the imaging mice to perform a sufficient number of trials on every session, the probe test and training problem sessions were modified so that rewards were available on two thirds of trials (instead of half) by doubling the number of reward available trails (on probe tests) and by increasing the number of rich tone trials (on training sessions). Additionally, the delay times between minimum wait and reward delivery were shortened by approximately 2s on average (distribution min = 0s, max = 54.63s, mean = 4.22s, median = 1.49s).

### Histology

Lens placement was determined post-mortem. Mice were transcardially perfused with 4% paraformaldehyde in phosphate buffered saline. Brains were removed and stored for at least 24 h in 4% paraformaldehyde before being transferred to a 30% sucrose solution for storage until they could be sliced at 50 μm, DAPI stained, and examined.

### Data analysis

#### Subject-level comparisons

We compared behavioral responses and reactivation scores using standard approaches. We used t-tests to evaluate differences between two groups at a single time point, between one group at two time points, or to compare one group at a single time point to zero. We used one-way repeated measures ANOVA tests to evaluate the inequality of one group at multiple time points, and mixed ANOVA tests to evaluate the interaction between two groups at multiple time points. In cases where the groups were unbalanced, such as in the imaging conditions, we used linear mixed effects models in place of mixed ANOVA tests. All statistics were performed in MATLAB (MathWorks, Natick, MA).

#### Modeling probe test responses

We fit a model to probe wait times for every subject to estimate their preferences for every tone. This allowed us to estimate preferences similarly for all subjects even if a subject did not provide data about their preference for a particular tone of interest (for example, if they did not run enough trials to sample all tones). To do this we used the *nlinfit* function in MATLAB to fit a curve to the wait times of each subject on each probe. We chose a linear combination of a normal and a logistic function (**Supplemental figure 1H**) as these described the various curve shapes observed in this study (**Figure 2A-B**). To ensure that we fit the most parsimonious model to each session, we first fit four separate models (intercept-only, intercept + logistic, intercept + normal, and intercept + logistic + normal) and then compared them by iteratively withholding a single data point from a session, fitting each model to the remaining wait times, and then measuring the error between each model and the withheld data point. After computing the error rates of each model, we selected the model with the lowest mean error rate as follows: we first selected a partial model by comparing the intercept + normal vs intercept + logistic models and kept the one with the lower error rate; we then compared the partial model vs the intercept-only model, and if the intercept-only model had the lower error rate we selected the intercept-only model; however, if the partial model had the lower error rate, then we compared the partial model vs the full (intercept + logistic + normal) model and selected the model with the lower error rate. In this way, we only selected the full model if it was more effective at predicting withheld data than any of the simpler models.

The intercept model was simply the mean wait time response in seconds. The intercept + logistic model was defined as

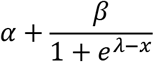

where *α* is the intercept, *β* is the height of the curve (in seconds), *λ* is the midpoint of the curve (the tone number, any value from 1 to 41). The intercept + normal model was written to constrain the width of the curve (in number of tones) so that it was proportionate to the height, defined as

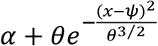

where *α* is the intercept, *θ* is the height of the curve and is proportional to the width of the curve, and *ψ* is the midpoint of the curve. The intercept + logistic + normal model was defined as

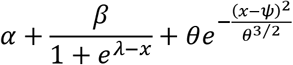

where *α* is the intercept, *β* is the height of the logistic curve, *λ* is the midpoint of the logistic curve, *θ* is the height and determines the width of the normal curve, and *ψ* is the midpoint of the normal curve.

#### Preprocessing calcium imaging data

Each imaging session produced as many videos as there were trials, along with the timestamps for each frame. We concatenated the imaging videos into a single session video using custom Python software, and then processed the resulting session video using CNMFe (Zhou et al., 2018) to identify the spatial footprints of, and to extract calcium signals from, individual neurons. The resulting calcium signals were then linearly up sampled to 100 Hz to match the sampling rate of the behavioral hardware.

#### Cell registration across sessions

In order to determine which neurons were active in each session, we used CellReg (Sheintuch et al., 2017) to align and register neuronal spatial footprints across all 19 imaging sessions for each mouse. Gross rotational differences between session videos, identified by comparing vasculature observable in maximum projection images, were manually corrected using custom rotation software written in MATLAB. We then registered the resulting footprints using rigid rotation and transformation in CellReg (**Supplemental figure 3**).

#### Time warping fluorescence signals

We obtained calcium signals from each trial for the series of events starting with nose poke and ending either when the mouse left the food cup or 2 s after reward delivery. To correct for slight differences in the time between each event on different trials, we linearly interpolated the observed calcium signals from each trial into a universal trial timeline with fixed durations between each event. The fixed durations (as seen in **Figure 5**) were approximately the median observed time between each event.

#### Mean activity correlations

We determined the similarity of neuronal activity patterns across pairs of sessions by first finding the mean time-warped activity pattern for each cell across all trials within a session and then computing the correlation between the mean activity patterns of a neuron in one session and the mean activity pattern of that same neuron in a second session. This analysis only included the portion of each trial that was common to both rewarded and unrewarded trials. The included events were the nose poke exit, the head entry into the food cup, the moment the tone on came on, and the end of the minimum wait time (2 s after tone on). Lastly, we computed the mean activity correlation between two sessions as the average correlation among all of the neurons that were active in both sessions.

#### Reactivation scores

We measured the reactivation of hippocampal ensemble activity in terms of the proportion of neurons that were active in two different sessions (proportion overlap) and how similar the activity of each neuron was across the two sessions (mean activity correlation). To do this, we used a reactivation score computed as

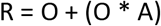

where R is the reactivation score, O is the proportion overlap, and A is the mean activity correlation. A score of 0 will occur if there is no overlap between the two sessions (no neurons are active in both sessions) or if all of the overlapping neurons have perfectly anticorrelated activity patterns between the two sessions. A score of 2 will only occur if there is perfect overlap (the exact same population of neurons is active in both sessions) and all of the neurons have perfectly correlated activity patterns.

#### Correcting for time between imaging sessions

Because the Structured training group learned new problems faster than the Unstructured training group (**Supplemental figure 2D-F**), there were slight differences in the number of days between the corresponding imaging sessions of each group (**Supplemental figure 4A**). We controlled for this in all of our measures of population similarity (population overlap, mean activity correlations, and reactivation scores) with the same general approach. We first computed the population similarity measure between every pair of imaging sessions within each Unstructured training group mouse and then pooled all of these values together. We only used Unstructured training sessions because these sessions showed a clear temporal trend that was unadulterated by learning related effects, and because the Structured training group did not have any long (> 50 days) durations between imaging sessions. However, similar results were obtained if we used sessions from both groups. We then estimated the effect of days between imaging sessions by fitting a truncated normal curve to the scatterplot of population similarity measurements using the *nlinfit* function in MATLAB with parameters for the intercept, height, width, and mid-point of the curve. For the mean activity correlation model, we assumed an intercept of 0 (no correlation between activity states as the time between imaging sessions approaches infinity) and therefore only fit parameters for height, width, and mid-point. Fit curves can be seen for population overlap and mean activity correlations in **Supplemental figures 4B-C** and for reactivation scores in **Supplemental figure 5A**. We then computed the new (time-corrected) population similarity measurements as the original measurement minus the value of the fit normal curve at the corresponding x-value (days between imaging sessions). This resulted in both positive and negative values, with negative values corresponding to population similarity below what was expected by the number of days between the two imaging sessions.

## References

Baraduc, P., Duhamel, J.-R., and Wirth, S. (2019). Schema cells in the macaque hippocampus. Science 363, 635–639. https://doi.org/10.1126/science.aav5404.

Barron, H.C., Reeve, H.M., Koolschijn, R.S., Perestenko, P.V., Shpektor, A., Nili, H., Rothaermel, R., Campo-Urriza, N., O’Reilly, J.X., Bannerman, D.M., et al. (2020). Neuronal Computation Underlying Inferential Reasoning in Humans and Mice. Cell 183, 228–243.e21. https://doi.org/10.1016/j.cell.2020.08.035.

Bartlett, F.C. (1932). Remembering: A study in experimental and social psychology (New York, NY, US: Cambridge University Press).

Behrens, T.E.J., Muller, T.H., Whittington, J.C.R., Mark, S., Baram, A.B., Stachenfeld, K.L., and Kurth-Nelson, Z. (2018). What Is a Cognitive Map? Organizing Knowledge for Flexible Behavior. Neuron 100, 490–509. https://doi.org/10.1016/j.neuron.2018.10.002.

Björk, E., Nevalainen, T., Hakumäki, M., and Voipio, H.M. (2000). R-weighting provides better estimation for rat hearing sensitivity. Lab Anim 34, 136–144. https://doi.org/10.1258/002367700780457518.

Bransford, J.D., and Johnson, M.K. (1972). Contextual prerequisites for understanding: Some investigations of comprehension and recall. Journal of Verbal Learning and Verbal Behavior 11, 717–726.

Brunec, I.K., and Momennejad, I. (2022). Predictive Representations in Hippocampal and Prefrontal Hierarchies. J Neurosci 42, 299–312. https://doi.org/10.1523/JNEUROSCI.1327-21.2021.

Bulkin, D.A., Law, L.M., and Smith, D.M. (2016). Placing memories in context: Hippocampal representations promote retrieval of appropriate memories. Hippocampus 26, 958–971. https://doi.org/10.1002/hipo.22579.

Butterly, D.A., Petroccione, M.A., and Smith, D.M. (2012). Hippocampal context processing is critical for interference free recall of odor memories in rats. Hippocampus 22, 906–913. https://doi.org/10.1002/hipo.20953.

Cai, D.J., Aharoni, D., Shuman, T., Shobe, J., Biane, J., Song, W., Wei, B., Veshkini, M., La-Vu, M., Lou, J., et al. (2016). A shared neural ensemble links distinct contextual memories encoded close in time. Nature 534, 115–118. https://doi.org/10.1038/nature17955.

Chase, W.G., and Simon, H.A. (1973). Perception in chess. Cognitive Psychology 4, 55–81. https://doi.org/10.1016/0010-0285(73)90004-2.

Chowdhury, A., Luchetti, A., Fernandes, G., Filho, D.A., Kastellakis, G., Tzilivaki, A., Ramirez, E.M., Tran, M.Y., Poirazi, P., and Silva, A.J. (2022). A locus coeruleus-dorsal CA1 dopaminergic circuit modulates memory linking. Neuron S0896-6273(22)00707-3. https://doi.org/10.1016/j.neuron.2022.08.001.

Craik, F.I.M., and Tulving, E. (1975). Depth of processing and the retention of words in episodic memory. Journal of Experimental Psychology: General 104, 268–294. https://doi.org/10.1037/0096-3445.104.3.268.

Dana, H., Chen, T.-W., Hu, A., Shields, B.C., Guo, C., Looger, L.L., Kim, D.S., and Svoboda, K. (2014). Thy1-GCaMP6 Transgenic Mice for Neuronal Population Imaging In Vivo. PLOS ONE 9, e108697. https://doi.org/10.1371/journal.pone.0108697.

Davis, H.P., and Squire, L.R. (1984). Protein synthesis and memory: a review. Psychol Bull 96, 518–559..

Debiec, J., LeDoux, J.E., and Nader, K. (2002). Cellular and systems reconsolidation in the hippocampus. Neuron 36, 527–538. https://doi.org/10.1016/s0896-6273(02)01001-2.

Dragoi, G., and Tonegawa, S. (2011). Preplay of future place cell sequences by hippocampal cellular assemblies. Nature 469, 397–401. https://doi.org/10.1038/nature09633.

Epsztein, J., Brecht, M., and Lee, A.K. (2011). Intracellular determinants of hippocampal CA1 place and silent cell activity in a novel environment. Neuron 70, 109–120. https://doi.org/10.1016/j.neuron.2011.03.006.

Fernández, R.S., Boccia, M.M., and Pedreira, M.E. (2016). The fate of memory: Reconsolidation and the case of Prediction Error. Neurosci Biobehav Rev 68, 423–441. https://doi.org/10.1016/j.neubiorev.2016.06.004.

Fuhs, M.C., and Touretzky, D.S. (2007). Context learning in the rodent hippocampus. Neural Comput 19, 3173–3215. https://doi.org/10.1162/neco.2007.19.12.3173.

Gershman, S.J., and Niv, Y. (2010). Learning latent structure: carving nature at its joints. Curr Opin Neurobiol 20, 251–256. https://doi.org/10.1016/j.conb.2010.02.008.

Gill, P.R., Mizumori, S.J.Y., and Smith, D.M. (2011). Hippocampal episode fields develop with learning. Hippocampus 21, 1240–1249. https://doi.org/10.1002/hipo.20832.

Girardeau, G., Benchenane, K., Wiener, S.I., Buzsáki, G., and Zugaro, M.B. (2009). Selective suppression of hippocampal ripples impairs spatial memory. Nat Neurosci 12, 1222–1223. https://doi.org/10.1038/nn.2384.

Gisquet-Verrier, P., and Riccio, D.C. (2018). Memory integration: An alternative to the consolidation/reconsolidation hypothesis. Prog Neurobiol 171, 15–31. https://doi.org/10.1016/j.pneurobio.2018.10.002.

Goshen, I., Brodsky, M., Prakash, R., Wallace, J., Gradinaru, V., Ramakrishnan, C., and Deisseroth, K. (2011). Dynamics of retrieval strategies for remote memories. Cell 147, 678–689. https://doi.org/10.1016/j.cell.2011.09.033.

Harlow, H.F. (1949). The formation of learning sets. Psychological Review 56, 51–65. https://doi.org/10.1037/h0062474.

Jacob, A.D., Ramsaran, A.I., Mocle, A.J., Tran, L.M., Yan, C., Frankland, P.W., and Josselyn, S.A. (2018). A Compact Head-Mounted Endoscope for In Vivo Calcium Imaging in Freely Behaving Mice. Curr Protoc Neurosci 84, e51. https://doi.org/10.1002/cpns.51.

Kelemen, E., and Fenton, A.A. (2010). Dynamic grouping of hippocampal neural activity during cognitive control of two spatial frames. PLoS Biol 8, e1000403. https://doi.org/10.1371/journal.pbio.1000403.

van Kesteren, M.T.R., Ruiter, D.J., Fernández, G., and Henson, R.N. (2012). How schema and novelty augment memory formation. Trends Neurosci 35, 211–219. https://doi.org/10.1016/j.tins.2012.02.001.

Knudsen, E.B., and Wallis, J.D. (2021). Hippocampal neurons construct a map of an abstract value space. Cell 184, 4640–4650.e10. https://doi.org/10.1016/j.cell.2021.07.010.

Kumaran, D., Hassabis, D., and McClelland, J.L. (2016). What Learning Systems do Intelligent Agents Need? Complementary Learning Systems Theory Updated. Trends Cogn Sci 20, 512–534. https://doi.org/10.1016/j.tics.2016.05.004.

Lee, J.L.C. (2009). Reconsolidation: maintaining memory relevance. Trends Neurosci 32, 413–420. https://doi.org/10.1016/j.tins.2009.05.002.

Lee, J.L.C., Nader, K., and Schiller, D. (2017). An Update on Memory Reconsolidation Updating. Trends Cogn Sci 21, 531–545. https://doi.org/10.1016/j.tics.2017.04.006.

Liu, X., Ramirez, S., Pang, P.T., Puryear, C.B., Govindarajan, A., Deisseroth, K., and Tonegawa, S. (2012). Optogenetic stimulation of a hippocampal engram activates fear memory recall. Nature 484, 381–385. https://doi.org/10.1038/nature11028.

Mack, M.L., Love, B.C., and Preston, A.R. (2018). Building concepts one episode at a time: The hippocampus and concept formation. Neurosci Lett 680, 31–38. https://doi.org/10.1016/j.neulet.2017.07.061.

Makino, H., and Komiyama, T. (2015). Learning enhances the relative impact of top-down processing in the visual cortex. Nat Neurosci 18, 1116–1122. https://doi.org/10.1038/nn.4061.

Markus, E.J., Qin, Y.L., Leonard, B., Skaggs, W.E., McNaughton, B.L., and Barnes, C.A. (1995). Interactions between location and task affect the spatial and directional firing of hippocampal neurons. J. Neurosci. 15, 7079–7094. https://doi.org/10.1523/JNEUROSCI.15-11-07079.1995.

Mau, W., Hasselmo, M.E., and Cai, D.J. (2020). The brain in motion: How ensemble fluidity drives memory-updating and flexibility. Elife 9, e63550. https://doi.org/10.7554/eLife.63550.

McCloskey, M., and Cohen, N.J. (1989). Catastrophic Interference in Connectionist Networks: The Sequential Learning Problem. In Psychology of Learning and Motivation, G.H. Bower, ed. (Academic Press), pp. 109–165.

McKenzie, S., Frank, A.J., Kinsky, N.R., Porter, B., Rivière, P.D., and Eichenbaum, H. (2014). Hippocampal representation of related and opposing memories develop within distinct, hierarchically organized neural schemas. Neuron 83, 202–215. https://doi.org/10.1016/j.neuron.2014.05.019.

McKenzie, S., Huszár, R., English, D.F., Kim, K., Christensen, F., Yoon, E., and Buzsáki, G. (2021). Preexisting hippocampal network dynamics constrain optogenetically induced place fields. Neuron 109, 1040–1054.e7. https://doi.org/10.1016/j.neuron.2021.01.011.

Miller, A.M.P., Mau, W., and Smith, D.M. (2019). Retrosplenial Cortical Representations of Space and Future Goal Locations Develop with Learning. Curr Biol 29, 2083–2090.e4. https://doi.org/10.1016/j.cub.2019.05.034.

Momennejad, I. (2020). Learning Structures: Predictive Representations, Replay, and Generalization. Curr Opin Behav Sci 32, 155–166. https://doi.org/10.1016/j.cobeha.2020.02.017.

Morton, N.W., and Preston, A.R. (2021). Concept formation as a computational cognitive process. Curr Opin Behav Sci 38, 83–89. https://doi.org/10.1016/j.cobeha.2020.12.005.

Muller, R.U., and Kubie, J.L. (1987). The effects of changes in the environment on the spatial firing of hippocampal complex-spike cells. J. Neurosci. 7, 1951–1968. https://doi.org/10.1523/JNEUROSCI.07-07-01951.1987.

Nieh, E.H., Schottdorf, M., Freeman, N.W., Low, R.J., Lewallen, S., Koay, S.A., Pinto, L., Gauthier, J.L., Brody, C.D., and Tank, D.W. (2021). Geometry of abstract learned knowledge in the hippocampus. Nature 595, 80–84. https://doi.org/10.1038/s41586-021-03652-7.

Niv, Y. (2019). Learning task-state representations. Nat Neurosci 22, 1544–1553. https://doi.org/10.1038/s41593-019-0470-8.

Park, A.J., Harris, A.Z., Martyniuk, K.M., Chang, C.-Y., Abbas, A.I., Lowes, D.C., Kellendonk, C., Gogos, J.A., and Gordon, J.A. (2021). Reset of hippocampal–prefrontal circuitry facilitates learning. Nature 591, 615–619. https://doi.org/10.1038/s41586-021-03272-1.

Payne, J.D., Schacter, D.L., Propper, R.E., Huang, L.-W., Wamsley, E.J., Tucker, M.A., Walker, M.P., and Stickgold, R. (2009). The role of sleep in false memory formation. Neurobiology of Learning and Memory 92, 327–334. https://doi.org/10.1016/j.nlm.2009.03.007.

Plitt, M.H., and Giocomo, L.M. (2021). Experience-dependent contextual codes in the hippocampus. Nat Neurosci 24, 705–714. https://doi.org/10.1038/s41593-021-00816-6.

Pudhiyidath, A., Morton, N.W., Viveros Duran, R., Schapiro, A.C., Momennejad, I., Hinojosa-Rowland, D.M., Molitor, R.J., and Preston, A.R. (2022). Representations of Temporal Community Structure in Hippocampus and Precuneus Predict Inductive Reasoning Decisions. Journal of Cognitive Neuroscience 34, 1736–1760. https://doi.org/10.1162/jocn_a_01864.

Purtle, R.B. (1973). Peak shift: A review. Psychological Bulletin 80, 408–421. https://doi.org/10.1037/h0035233.

Rashid, A.J., Yan, C., Mercaldo, V., Hsiang, H.-L.L., Park, S., Cole, C.J., De Cristofaro, A., Yu, J., Ramakrishnan, C., Lee, S.Y., et al. (2016). Competition between engrams influences fear memory formation and recall. Science 353, 383–387. https://doi.org/10.1126/science.aaf0594.

Redish, A.D., Jensen, S., Johnson, A., and Kurth-Nelson, Z. (2007). Reconciling reinforcement learning models with behavioral extinction and renewal: implications for addiction, relapse, and problem gambling. Psychol Rev 114, 784–805. https://doi.org/10.1037/0033-295X.114.3.784.

Richards, B.A., Xia, F., Santoro, A., Husse, J., Woodin, M.A., Josselyn, S.A., and Frankland, P.W. (2014). Patterns across multiple memories are identified over time. Nat Neurosci 17, 981–986. https://doi.org/10.1038/nn.3736.

Robinson, N.T.M., Descamps, L.A.L., Russell, L.E., Buchholz, M.O., Bicknell, B.A., Antonov, G.K., Lau, J.Y.N., Nutbrown, R., Schmidt-Hieber, C., and Häusser, M. (2020). Targeted Activation of Hippocampal Place Cells Drives Memory-Guided Spatial Behavior. Cell 183, 1586–1599.e10. https://doi.org/10.1016/j.cell.2020.09.061.

Sadtler, P.T., Quick, K.M., Golub, M.D., Chase, S.M., Ryu, S.I., Tyler-Kabara, E.C., Yu, B.M., and Batista, A.P. (2014). Neural constraints on learning. Nature 512, 423–426. https://doi.org/10.1038/nature13665.

Samborska, V., Butler, J., Walton, M., Behrens, T.E.J., and Akam, T. (2021). Complementary task representations in hippocampus and prefrontal cortex for generalising the structure of problems. 2021.03.05.433967. https://doi.org/10.1101/2021.03.05.433967.

Sanders, H., Wilson, M.A., and Gershman, S.J. (2020). Hippocampal remapping as hidden state inference. Elife 9, e51140. https://doi.org/10.7554/eLife.51140.

Schapiro, A.C., Turk-Browne, N.B., Norman, K.A., and Botvinick, M.M. (2016). Statistical learning of temporal community structure in the hippocampus. Hippocampus 26, 3–8. https://doi.org/10.1002/hipo.22523.

Schlichting, M.L., and Frankland, P.W. (2017). Memory allocation and integration in rodents and humans. Current Opinion in Behavioral Sciences 17, 90–98. https://doi.org/10.1016/j.cobeha.2017.07.013.

Sheintuch, L., Rubin, A., Brande-Eilat, N., Geva, N., Sadeh, N., Pinchasof, O., and Ziv, Y. (2017). Tracking the Same Neurons across Multiple Days in Ca2+ Imaging Data. Cell Rep 21, 1102–1115. https://doi.org/10.1016/j.celrep.2017.10.013.

Smith, D.M., and Mizumori, S.J.Y. (2006). Learning-related development of context-specific neuronal responses to places and events: the hippocampal role in context processing. J Neurosci 26, 3154–3163. https://doi.org/10.1523/JNEUROSCI.3234-05.2006.

Smith, S.M., and Vela, E. (2001). Environmental context-dependent memory: A review and meta-analysis. Psychonomic Bulletin & Review 8, 203–220. https://doi.org/10.3758/BF03196157.

Squire, L.R. (2004). Memory systems of the brain: A brief history and current perspective. Neurobiology of Learning and Memory 82, 171–177. https://doi.org/10.1016/j.nlm.2004.06.005.

Sun, C., Yang, W., Martin, J., and Tonegawa, S. (2020). Hippocampal neurons represent events as transferable units of experience. Nat Neurosci 23, 651–663. https://doi.org/10.1038/s41593-020-0614-x.

Takehara-Nishiuchi, K., and McNaughton, B.L. (2008). Spontaneous changes of neocortical code for associative memory during consolidation. Science 322, 960–963. https://doi.org/10.1126/science.1161299.

Tanaka, K.Z., Pevzner, A., Hamidi, A.B., Nakazawa, Y., Graham, J., and Wiltgen, B.J. (2014). Cortical Representations Are Reinstated by the Hippocampus during Memory Retrieval. Neuron 84, 347–354. https://doi.org/10.1016/j.neuron.2014.09.037.

Tervo, D.G.R., Tenenbaum, J.B., and Gershman, S.J. (2016). Toward the neural implementation of structure learning. Curr Opin Neurobiol 37, 99–105. https://doi.org/10.1016/j.conb.2016.01.014.

Tolman, E.C. (1948). Cognitive maps in rats and men. Psychol Rev 55, 189–208. https://doi.org/10.1037/h0061626.

Tse, D., Langston, R.F., Kakeyama, M., Bethus, I., Spooner, P.A., Wood, E.R., Witter, M.P., and Morris, R.G.M. (2007). Schemas and memory consolidation. Science 316, 76–82. https://doi.org/10.1126/science.1135935.

Tse, D., Takeuchi, T., Kakeyama, M., Kajii, Y., Okuno, H., Tohyama, C., Bito, H., and Morris, R.G.M. (2011). Schema-Dependent Gene Activation and Memory Encoding in Neocortex. Science 333, 891–895. https://doi.org/10.1126/science.1205274.

Vikbladh, O.M., Meager, M.R., King, J., Blackmon, K., Devinsky, O., Shohamy, D., Burgess, N., and Daw, N.D. (2019). Hippocampal Contributions to Model-Based Planning and Spatial Memory. Neuron 102, 683–693.e4. https://doi.org/10.1016/j.neuron.2019.02.014.

Whittington, J.C.R., Muller, T.H., Mark, S., Barry, C., and Behrens, T.E.J. (2018). Generalisation of structural knowledge in the hippocampal-entorhinal system. https://doi.org/10.48550/arXiv.1805.09042.

Whittington, J.C.R., Muller, T.H., Mark, S., Chen, G., Barry, C., Burgess, N., and Behrens, T.E.J. (2020). The Tolman-Eichenbaum Machine: Unifying Space and Relational Memory through Generalization in the Hippocampal Formation. Cell 183, 1249–1263.e23. https://doi.org/10.1016/j.cell.2020.10.024.

Whittington, J.C.R., McCaffary, D., Bakermans, J.J.W., and Behrens, T.E.J. (2022). How to build a cognitive map: insights from models of the hippocampal formation. https://doi.org/10.48550/arXiv.2202.01682.

Wiltgen, B.J., and Silva, A.J. (2007). Memory for context becomes less specific with time. Learn. Mem. 14, 313–317. https://doi.org/10.1101/lm.430907.

Wood, E.R., Dudchenko, P.A., Robitsek, R.J., and Eichenbaum, H. (2000). Hippocampal neurons encode information about different types of memory episodes occurring in the same location. Neuron 27, 623–633. https://doi.org/10.1016/s0896-6273(00)00071-4.

Zeithamova, D., Dominick, A.L., and Preston, A.R. (2012). Hippocampal and ventral medial prefrontal activation during retrieval-mediated learning supports novel inference. Neuron 75, 168–179. https://doi.org/10.1016/j.neuron.2012.05.010.

Zhou, P., Resendez, S.L., Rodriguez-Romaguera, J., Jimenez, J.C., Neufeld, S.Q., Giovannucci, A., Friedrich, J., Pnevmatikakis, E.A., Stuber, G.D., Hen, R., et al. (2018). Efficient and accurate extraction of in vivo calcium signals from microendoscopic video data. Elife 7, e28728. https://doi.org/10.7554/eLife.28728.

